# Social Disconnection in the Brain: Loneliness and Age across Networks using Graph Theory

**DOI:** 10.64898/2026.02.03.703621

**Authors:** Yen-Wen Chen, Turhan Canli

## Abstract

Loneliness, conceptualized as a multi-dimensional construct of unmet social needs, has been linked to adverse health outcomes across the lifespan, prompting significant interest in its underlying neural processes. Our study aimed to address the limitations of prior neuroimaging studies of loneliness by leveraging the Lifespan Human Connectome Project Aging dataset and applying graph theory to characterize its relationship with age and resting-state brain network organization. Socio-demographic measures confirmed prior work that higher loneliness was associated with younger age, being male, unmarried, and living alone. While loneliness showed no main effects on neural graph measures, a significant interaction between loneliness and age emerged for the local interconnectivity of the Default Model and Frontoparietal networks after adjusting for key socio-demographic factors. Conversely, older age was associated with lower functional connectivity, reduced global efficiency, and less modular brain network organization. Different graph measures showed distinct age-related associations, highlighting the heterogeneous nature of brain aging. The absence of a main effect of loneliness, while unexpected, underscores the complex, subjective nature of loneliness and suggests that its neural correlates may manifest differently across ages.

---

Humans are inherently social, and while social connectedness improve our chance of survival, losing social connections can be detrimental to it. Loneliness describes the subjective emotional experience of perceived social isolation, in which one’s social needs are not satisfied by the quantity and quality of one’s social relationships (Cacioppo, et al., 2006; Hawkley & Cacioppo, 2010). It is a multidimensional construct encompassing isolation, relational connectedness, and collective connectedness (Hawkley et al., 2005; Weiss, 1973).

Loneliness negatively impacts physical and mental health beyond objective social factors like social network size or social engagement (Cacioppo & Hawkley, 2009; Gow et al., 2007; R. S. Wilson et al., 2007). Across the lifespan, loneliness has been linked to poorer psychological well-being, mental disorders, and increased health risks (Beutel et al., 2017; Cacioppo et al., 2000, 2002, 2006, 2010; Ge et al., 2017; Hawkley et al., 2003, 2009, 2010; Mwilambwe-Tshilobo et al., 2019; Perissinotto et al., 2012; Pressman et al., 2005; Shankar et al., 2011; Vanhalst et al., 2013; Wang et al., 2023; Yi et al., 2018). Among older adults, it is associated with accelerated cognitive decline, increased risk of dementia and Alzheimer’s disease, and higher mortality (Donovan et al., 2017; Holt-Lunstad et al., 2015; Holwerda et al., 2016; Kim et al., 2021; Luo et al., 2012; R. S. Wilson et al., 2007). Although interventions to reduce loneliness have been developed, their effects remain modest and have not yet demonstrated consistent health benefits (Ellard et al., 2022; Hickin et al., 2021; Masi et al., 2011).

Loneliness is experienced across all ages and is shaped by diverse socio-demographic factors, including age, sex, marital status, living arrangement, and socioeconomic status (Beutel et al., 2017; Böger & Huxhold, 2018; Cacioppo et al., 2010, 2013; Distel et al., 2010; Dykstra et al., 2005; Ge et al., 2017; Hawkley et al., 2005, 2010; Luhmann & Hawkley, 2016; Perissinotto et al., 2012; Pinquart & Sorensen, 2001; Qualter et al., 2015; Routasalo et al., 2006; Savikko et al., 2005). These factors likely reflect age-specific social contexts and resources, underscoring the importance of considering socio-demographic variability when studying loneliness.

Neuroimaging research has begun to identify the underlying brain mechanisms of loneliness. Prior work links loneliness to altered gene expression in reward- and control-related regions (Canli et al., 2017, 2018), differences in brain activity during social and self-referential processing (Cacioppo et al., 2009; Courtney & Meyer, 2020; D’Agostino et al., 2018; Golde et al., 2019; Inagaki et al., 2016), and structural variation in regions supporting mentalizing, emotion regulation, and executive control (Düzel et al., 2019; Kanai et al., 2012; Kiesow et al., 2020; Kong et al., 2015; Liu et al., 2016; Meng et al., 2017; Nakagawa et al., 2015; Tian et al., 2014). Emerging evidence suggests that loneliness moderates age–brain relationships, including steeper age-related amygdala volume loss (Düzel et al., 2019), increased brain age relative to chronological age (Lange et al., 2021), and age-related cerebral blood flow differences in social cognition regions (Chen et al., 2021). These findings suggest that loneliness may accelerate brain aging.

Resting-state functional connectivity (RSFC) provides a system-level approach to examine intrinsic brain organization (Fox et al., 2005; Fox & Raichle, 2007). Loneliness has been associated with reduced large-scale RSFC as well as altered within- and between-network connectivity across the brain (Feng et al., 2019; Layden et al., 2017; Mwilambwe-Tshilobo et al., 2019; Spreng et al., 2020; Tian et al., 2017). Beyond pairwise connectivity, graph-theoretical approaches model the brain as an integrated network and characterize its organizational properties (Andellini et al., 2015; Bullmore & Sporns, 2009; van den Heuvel & Hulshoff Pol, 2010). Given that loneliness is associated with widely distributed neural systems (Lam et al., 2021), graph-based analyses are well-suited for detecting network-level differences in brain organization associated with loneliness. Specifically, different graph metrics capture complementary aspects of network topology, including nodal centrality (e.g., degree and betweenness centrality), communication efficiency and integration (e.g., shortest path length), local segregation (e.g., clustering coefficient), inter-network connectivity (participation coefficient), and community structure (e.g., modularity) (Fornito et al., 2016). Because prior neuroimaging studies of loneliness have predominantly examined pairwise functional connectivity, we adopt an exploratory approach, examining whether loneliness is associated with these complementary graph-theoretic properties.

The current study leveraged large-scale data from the Lifespan Human Connectome Project (HCP) Aging (Bookheimer et al., 2019) to address gaps in the literature, which has predominantly focused on younger (ages 20–30) or older adults (ages 60+). By applying a graph-theoretical framework across a broad adult age range, we aimed to characterize how loneliness and its socio-demographic correlates interact with age to shape brain network organization.

## Methods

### Participants

Our study utilized data from the Lifespan HCP Aging 2.0 release dataset (Bookheimer et al., 2019). HCP-Aging recruited a typical aging population in the absence of cognitive impairment due to pathological causes and screened for cognitive ability. The 2.0 release dataset included 725 participants aged between 36–100 years old (*M_age_* = 60.36 ± 15.73), 406 females and 319 males. In this study, participants were further excluded if they (1) did not complete loneliness survey (*n* = 93), (2) did not complete four runs of resting-state fMRI scans (*n* = 6), (3) had any resting-state fMRI scan was excluded due to excessive motion (*n* = 130) or did not pass the data quality assessment (*n* = 1). The final sample in this study included 512 participants (*M_age_* = 58.23 ± 14.87, 303 females).

### Behavioral Measures

#### Socio-Demographic Factors

Socio-demographic information, including age, sex, marital status, household size, employment status, and income, was collected using Semi-Structured Assessment for the Genetics of Alcoholism–IV (Bucholz et al., 2015). Education was collected using MoCA.

#### NIH-Toolbox Loneliness Survey

Loneliness was assessed using NIH-Toolbox Loneliness Survey (Gershon et al., 2010). The Loneliness Survey assesses an individual’s perception of loneliness using five 5-point (*Never*–*Always*) items. A sample item is “In the past month, I feel alone and apart from others.” HCP-Aging provided uncorrected standard score (T-score), which compared the performance of the nationally representative normative sample. T-scores were used in the study.

#### Montreal Cognitive Assessment

General cognition was measured by Montreal Cognitive Assessment (MoCA) (Nasreddine et al., 2005). MoCA measures seven cognitive domains: visuospatial/executive, naming, attention, language, abstraction, delayed recall, and orientation with a maximum score of 30. HCP-A excluded participants if the total score was below 20 for ages 36–79 years, or below 18 for ages 80 and above. MoCA scores were corrected for education.

### Behavioral Data Preparation

#### Data Categorization and Transformation for Socio-Demographic Factors

**Age.** In years.
**Sex** Males and females.
**Marital Status.** Dichotomized into Married (including ‘married’ and ‘living as married’) and Unmarried (including ‘never married’, ‘divorced’, ‘separated’, and ‘widowed’).
**Household Status.** Dichotomized into Living Alone and Living with Others.
**Employment Status.** Employed and Unemployed.
**Income.** The total combined family income for the past 12 months. Due to the positively skewed data, income was transformed with natural logarithm.
**Education.** Education was categorized into three groups: High School and Below, College, and Graduate.

### MRI Data Acquisition and Preprocessing

MRI data were acquired on Siemens 3T Prisma scanner with 32-channel head coil (Harms et al., 2018). Structural T1-weighted and resting-state fMRI were collected using HCP-Aging acquisition sequences, including multi-echo T1w and multiband EPI resting-state scans, with participants instructed to fixate on a cross during rest. Detailed acquisition has been reported previously (Elam et al., 2021; Harms et al., 2018).

Data were processed using the HCP minimal preprocessing pipelines (Glasser et al., 2013; Smith et al., 2013). Structural and resting-state fMRI data underwent distortion correction, normalization, surface and volume mapping to CIFTI grayordinate space. Artifacts were removed using ICA-FIX non-aggressive denoising (Glasser et al., 2016; Robinson et al., 2014; Smith et al., 2013). Finally, cross-subject functional alignment was performed using Multi-modal Surface Matching algorithm (Glasser et al., 2016; Robinson et al., 2014). Detailed preprocessing procedures have been described in the primary HCP literature. Considering that resting-state functional images are sensitive to motion, participants with excessive high-motion frames were excluded in this study. High-motion frame was defined by framewise displacement (FD) > 0.5mm and the frame prior and after this frame were also marked as high-motion. Participants were excluded if any of the four runs had more than 40% of high-motion frames.

### Resting-State fMRI Functional Connectivity Matrix Construction

We used the Cole-Anticevic Brain Network Parcellation (CAB-NP, v1.1.6) (Ji et al., 2019), which consists of 360 cortical parcels and 358 subcortical parcels, to construct RSFC matrix. 718 parcels were clustered into 12 networks: Primary Visual, Secondary Visual, Somatomotor, Auditory, Default Mode, Cingulo-Opercular, Frontoparietal, Dorsal Attention, Orbito-Affective, Language, Ventral Multimodal, and Posterior Multimodal. Each resting scan was first de-meaned, variance normalized, and parcellated using Connectome Workbench v1.5.0 (Marcus et al., 2013). A full correlation was computed and Fisher’s z transformed for each resting scan. Finally, four functional connectivity matrices were averaged for each participant. The code for functional connectivity matrix construction is available on https://osf.io/p6srv/.

### Resting-State Functional Connectivity Graph Theoretical Based Measures

The brain functional organization was modeled as a graph of nodes (brain parcels) and edges (functional connections) using an undirected, weighted RSFC matrix. Given ongoing debating regarding the biological interpretation of the negative connectivity weights (Chang & Glover, 2009; G. Chen et al., 2011; Fox et al., 2009; Murphy et al., 2009), we utilized the absolute value of the correlation (Centeno et al., 2022). Prior work suggested this approach yielded comparable within- and between-network connectivity and segregation metrics to positive-only weighting (Chan et al., 2014). To assess robustness to edge-weight definition, supplementary analyses were also conducted using positive-only and negative-only connectivity matrices (see Supplementary Materials). Graph measures were computed using NetworkX 2.4 (Hagberg et al., 2008), with analysis code available on https://osf.io/p6srv/. Primary analyses focused on representative graph metrics spanning complementary aspects of network topology, including degree centrality, betweenness centrality, shortest path length, clustering coefficient, participation coefficient, and modularity. Additional related metrics (closeness centrality and eigenvector centrality) are reported in the Supplementary Materials to reduce redundancy among highly correlated graph measures.

#### Nodal Degree/Strength

Nodal degree (or strength) represents the sum of the connectivity weights of a node. A normalized measure of *nodal strength* was used by taking the average of the number of connections a node had (Fornito et al., 2016).

#### Shortest Path Length

Path length between two nodes, calculated as the sum of reciprocal weights, reflects the efficiency of information transfer. The *shortest path* was calculated with Dijkstra’s algorithm. The *Average Shortest Path Length* (or *Characteristic Path Length*) across all node pairs represents network integration efficiency, with shorter paths indicating more efficient information routing (Fornito et al., 2016).

#### Betweenness Centrality

*Betweenness centrality* measures how often a node lies on the shortest paths between other nodes, calculated as the sum of the fraction of all pairs of nodes shortest paths that pass through a node. Higher values indicate nodes that are critical for information flow (Fornito et al., 2016).

#### Clustering Coefficient

*Clustering coefficient* quantifies local interconnectivity, calculated as the geometric average of a node’s subgraph edge weights. Higher values indicate tightly connected local clusters that may support forming functional specialized clusters (Fornito et al., 2016).

#### Participation Coefficient

*Participation coefficient* quantifies the extent to which a node connects to other networks based on 12 CAB-NP network partitions, calculated as one minus the squared ratio of within-module degree to total degree (Guimerà & Amaral, 2005). Higher values indicate greater inter-network communication. Impairment of such nodes is associated with widespread influence across the whole brain (Fornito et al., 2016; Letina et al., 2019).

#### Modularity

*Modularity* is a global metric quantifying the extent to which network nodes form functional communities (Clauset et al., 2004; Newman & Girvan, 2004). Using the 12 CAB-NP partitions, the *modularity* index (*Q*) was calculated as the difference between observed within-community edges and those expected by chance. Higher value indicates greater functional specialization.

### Behavioral Statistical Analysis

Behavioral analyses were conducted in R v4.0.3 (Team R Core, 2017). The locally estimated scatterplot smoothing plots showed no evidence of non-linearity between loneliness and continuous variables (age, income), so these variables were treated linearly. Correlations between loneliness and socio-demographic factors were examined with Pearson *r* correlation or Analysis of Variance (ANOVA). Factors showing significant associations with loneliness were entered into a multiple linear regression to identify robust predictors and potential interactions. Significance was defined as *p*-value of < .05.

### Loneliness and Age – Graph-Based Brain Functional Connectivity Statistical Analysis

Association of loneliness, age, and their interaction with graph measures was estimated using linear regression, separately for each graph measure. Significance was assessed using non-parametric permutation tests (5,000 iterations) implemented in FSL Permutation Analysis of Linear Models (PALM, v119-alpha) (Winkler et al., 2014). All analyses were adjusted for sex, MoCA, mean FD, total gray matter volume, and scanning sites. Exchangeability blocks were used to estimate variance within each of four scanning sites. For each graph metric, family-wise error (FWE) correction was applied jointly across 718 parcels and three contrasts of interest (age, loneliness, and their interaction) using permutation-based inference in PALM. This correction was performed separately for each graph metric, treating each metric as an independent family of tests. Significant associations were defined as two-tailed FWE-corrected p-value < .05. Furthermore, to verify the multicollinearity did not inflate the interaction estimates, variance inflating factor (VIF) was computed for all predictors, including variables of no interest. All VIFs were below 1.9, indicating negligible multicollinearity.

## Results

### Loneliness and Socio-Demographic Associations

Bivariate analyses with loneliness showed that people who were younger, unmarried, and living alone reported higher levels of loneliness (Table 1, Figure 1). No differences in sex, employment status, or education were observed. Annual family income was negatively associated with loneliness. When all socio-demographic factors were included simultaneously, age, sex, marital status, and household status were robustly associated with loneliness (Table 2, *R_adj_^2^* = .14, *F*(8, 357) = 8.20, *p* = 3.35e-10). Younger individuals, males, unmarried status, and those living alone reported higher levels of loneliness.

**Figure 1.**
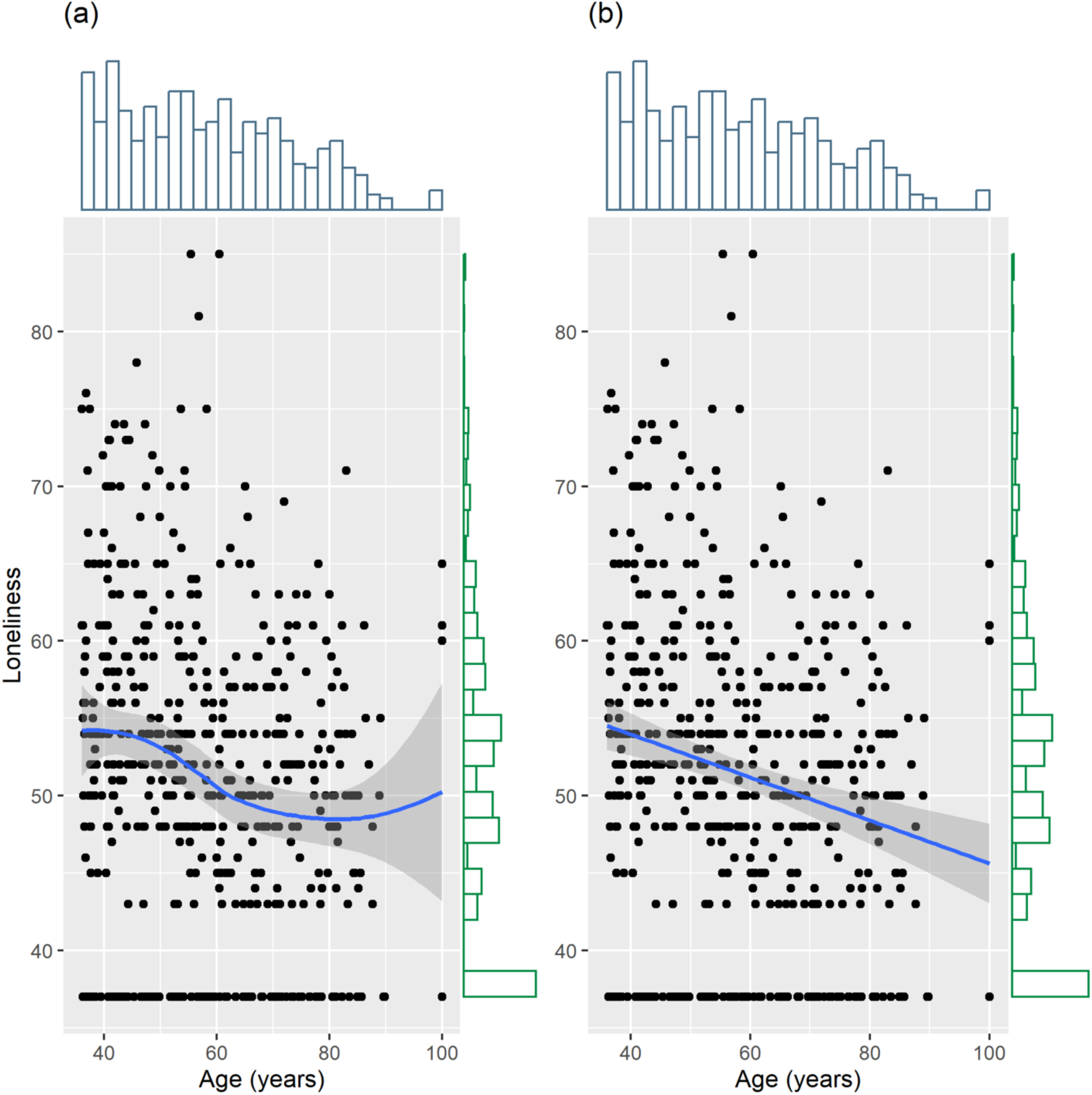
Distribution and Association of Loneliness and Age. (a) Association between loneliness and age with locally estimated scatterplot smoothing. (b) Association between loneliness and age with linear regression line.

**Table 1:**
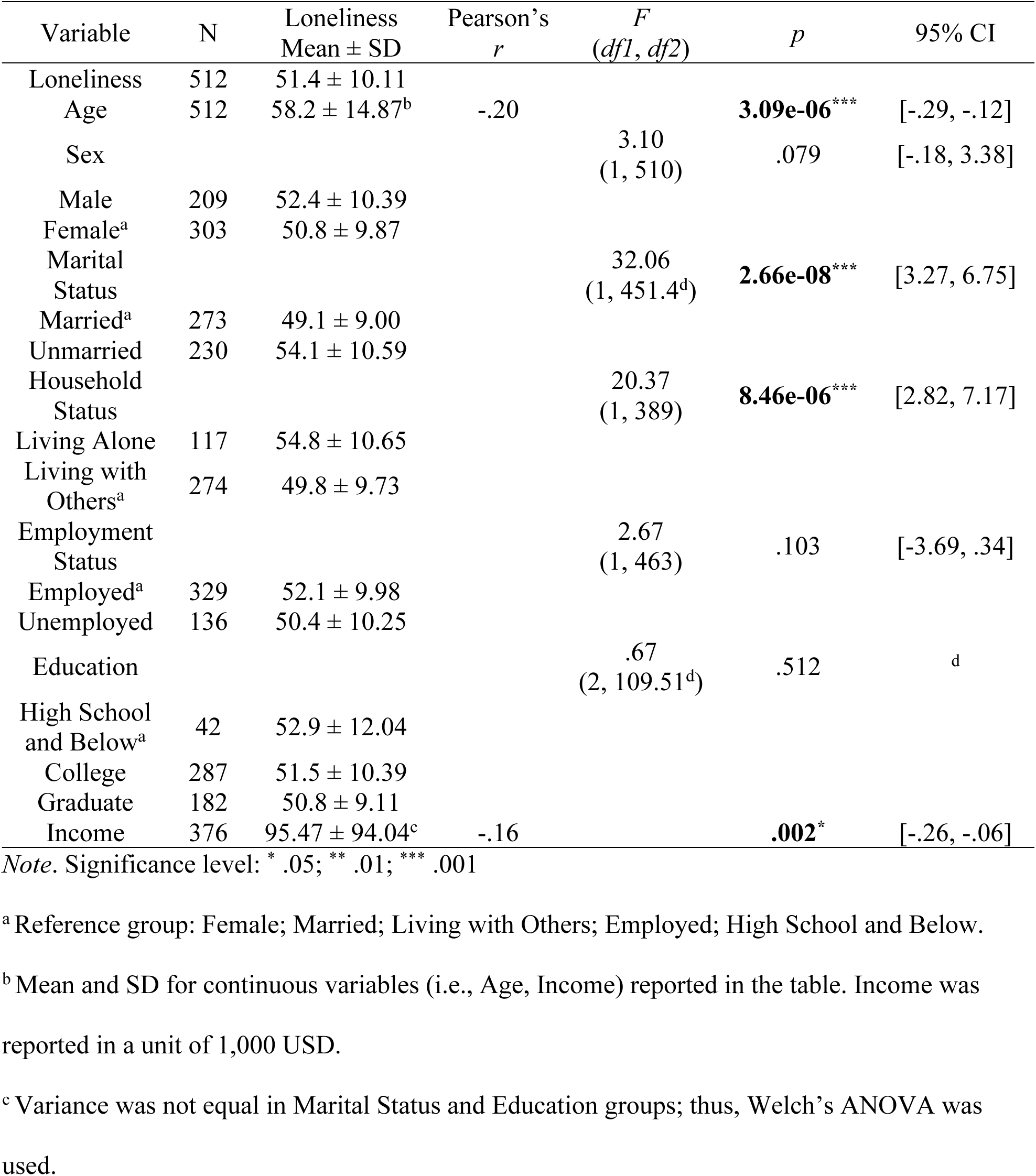
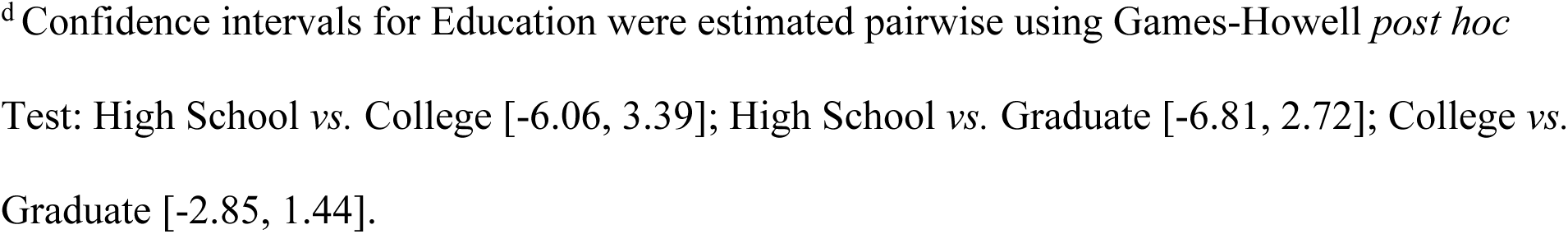
Participant Demographics and Behavioral Measures Descriptive Statistics and Association with Loneliness.

| Variable | N | Loneliness<br>Mean ± SD | Pearson's<br>*r* | *F*<br>(*df1*, *df2*) | *p* | 95% CI |
| --- | --- | --- | --- | --- | --- | --- |
| Loneliness | 512 | 51.4 ± 10.11 | | | | |
| Age | 512 | 58.2 ± 14.87<sup>b</sup> | -.20 | | **3.09e-06\*\*\*** | [-.29, -.12] |
| Sex |  |  |  | 3.10<br>(1, 510) | .079 | [-.18, 3.38] |
| Male | 209 | 52.4 ± 10.39 | | | | |
| Female<sup>a</sup> | 303 | 50.8 ± 9.87 | | | | |
| Marital<br>Status |  |  |  | 32.06<br>(1, 451.4<sup>d</sup>) | **2.66e-08\*\*\*** | [3.27, 6.75] |
| Married<sup>a</sup> | 273 | 49.1 ± 9.00 | | | | |
| Unmarried | 230 | 54.1 ± 10.59 | | | | |
| Household<br>Status |  |  |  | 20.37<br>(1, 389) | **8.46e-06\*\*\*** | [2.82, 7.17] |
| Living Alone | 117 | 54.8 ± 10.65 | | | | |
| Living with<br>Others<sup>a</sup> | 274 | 49.8 ± 9.73 | | | | |
| Employment<br>Status |  |  |  | 2.67<br>(1, 463) | .103 | [-3.69, .34] |
| Employed<sup>a</sup> | 329 | 52.1 ± 9.98 | | | | |
| Unemployed | 136 | 50.4 ± 10.25 | | | | |
| Education |  |  |  | .67<br>(2, 109.51<sup>d</sup>) | .512 | <sup>d</sup> |
| High School<br>and Below<sup>a</sup> | 42 | 52.9 ± 12.04 | | | | |
| College | 287 | 51.5 ± 10.39 | | | | |
| Graduate | 182 | 50.8 ± 9.11 | | | | |
| Income | 376 | 95.47 ± 94.04<sup>c</sup> | -.16 | | **.002\*** | [-.26, -.06] |

Note. Significance level: \* .05; \*\* .01; \*\*\* .001
<sup>a</sup> Reference group: Female; Married; Living with Others; Employed; High School and Below.
<sup>b</sup> Mean and SD for continuous variables (i.e., Age, Income) reported in the table. Income was reported in a unit of 1,000 USD.
<sup>c</sup> Variance was not equal in Marital Status and Education groups; thus, Welch's ANOVA was used.

**Table 2:**
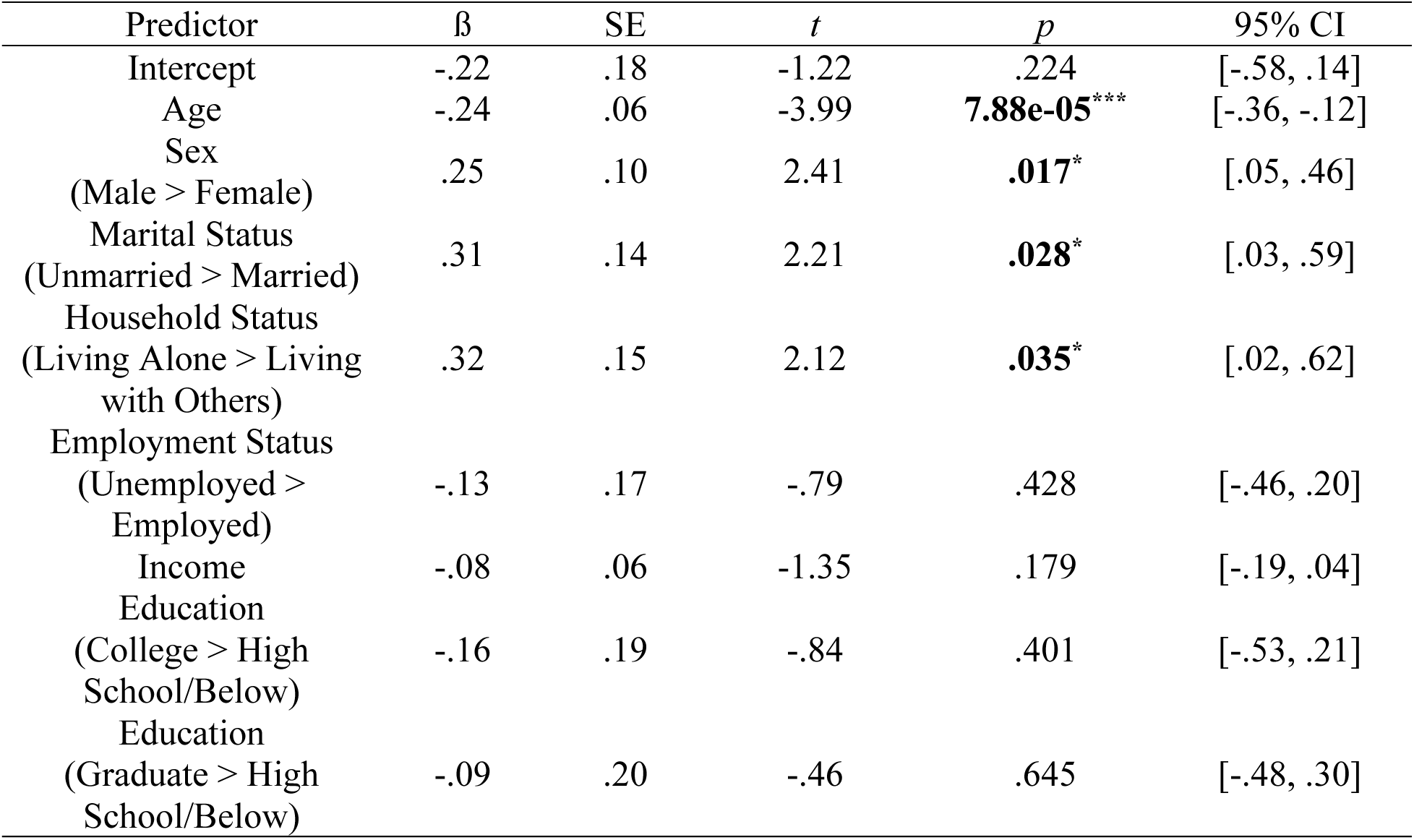
Regression Results of the Association between Loneliness and Socio-Demographic Factors.

### Loneliness and Graph-Based Brain Functional Connectivity Measures

To examine the association of loneliness and age with RSFC, loneliness, age, and their interaction were used as main predictors to examine their association with graph measures described in the Methods. Neither a significant main effect of loneliness, nor a significant interaction of loneliness and age, was revealed with any of the nodal or global graph measures. Supplementary analyses using positive-only and negative-only connectivity matrices yielded broadly consistent patterns with the main results. Specifically, results from the positive-only connectivity largely replicated findings from the absolute-weighted analyses, whereas negative-only connectivity showed substantially attenuated effects across metrics. No associations with loneliness or loneliness x age interaction were observed in either decomposition (see Supplementary Materials).

Given that behavioral results showed a significant association between socio-demographic factors (i.e., sex, marital status, and household status) and loneliness, we conducted a separate set of linear regressions that adjusted for those socio-demographic factors (in addition to MoCA, mean FD, total GMV, and scanning sites) for a subset of participants (*n* = 390) with available data. No significant main effect of loneliness was associated with any nodal or global graph measures. However, the interaction between loneliness and age was significantly associated with the *clustering coefficient* in two nodes: the left caudate in the Default Mode Network (*z* = −3.28, *p_FWE_* = .0156) and the right cerebellum in the Frontoparietal Network (*z* = −3.24, *p_FWE_* = .0202). To elucidate the results of the interaction, Figure 2–3 illustrate the loneliness x age interaction for each significance node by categorizing the continuous age variable into 10-year intervals; note that all statistical analyses retained age as a continuous variable and the age groupings are for descriptive visualization purpose only. For both nodes, the association with the *clustering coefficient* is positive for younger individuals (∼ under 65 years old), but negative for older individuals aged (∼ over 65 years old), reflecting an age-dependent influence on the association between loneliness and brain network organization.

**Figure 2.**
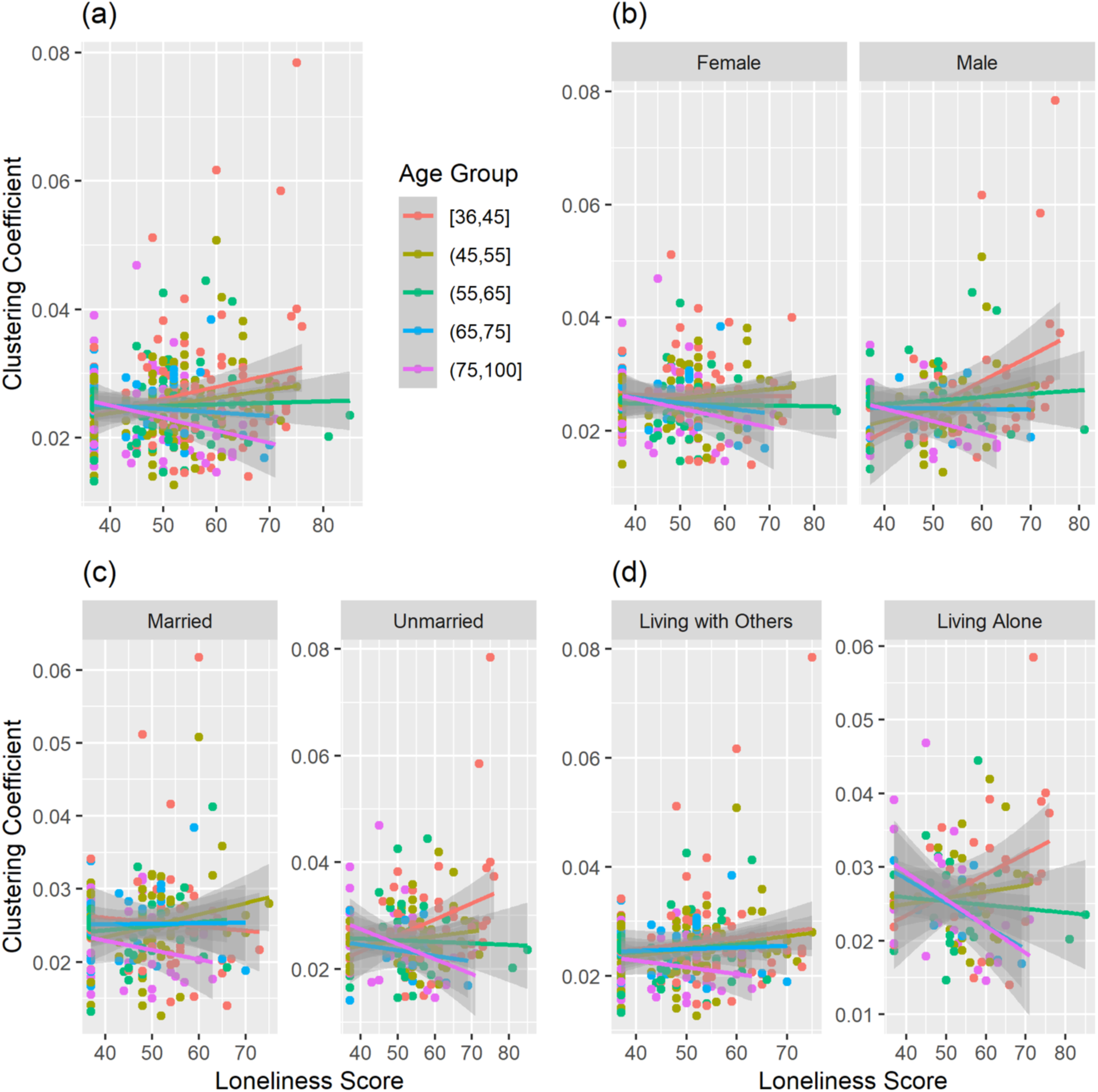
Interaction between Loneliness and Age on the Clustering Coefficient of Default Mode Network (Left Caudate). (a) 390 participants, (b) by sex, (c) by marital status, and (d) by household status.

**Figure 3.**
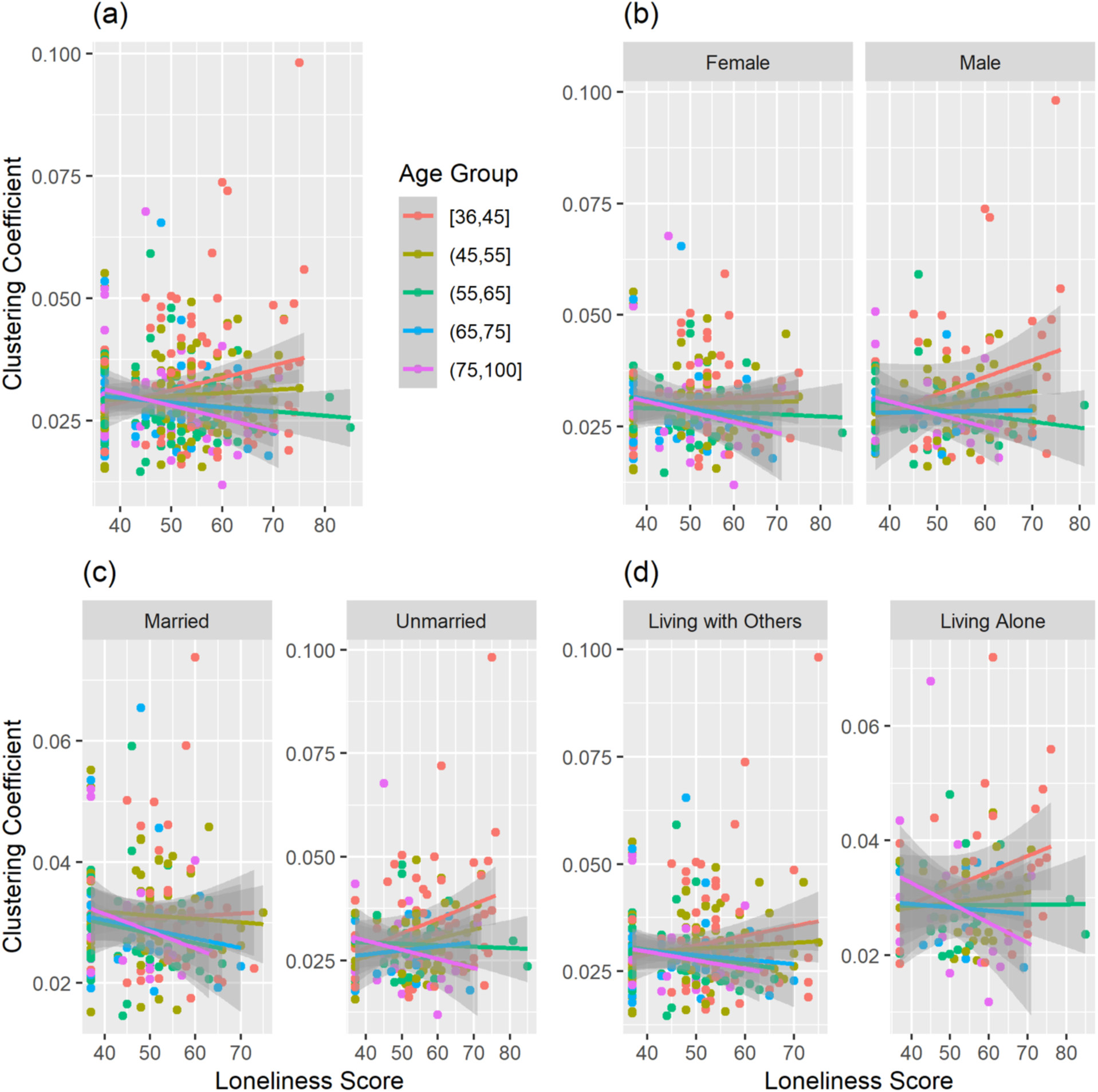
Interaction between Loneliness and Age on the Clustering Coefficient of Frontoparietal Network (Right Cerebellum). (a) 390 participants, (b) by sex, (c) by marital status, and (d) by household status.

### Age and Graph-Based Brain Functional Connectivity Measures

Whereas there was no main effect of loneliness, there was a significant main effect of age associated with all graph measures. Table 3 summarizes the age results at the network level and Figure 4 showed the un-thresholded association of age and nodal graph measures. Table S1–S6 lists the results of age and nodal graph measures. *Nodal Strength*: The sum of the connectivity weights of a node. Age was negatively associated with *normalized strength* of nodes across 10 networks defined by CAB-NP (Ji et al., 2019), particularly in the Secondary Visual Network and Cingulo-Opercular Network (Detailed nodal results are provided in Supplementary Table S1). No positive age-related association was found. *Betweenness Centrality*: A measure of how often a node is on the shortest path between two other nodes. Age was positively associated with *betweenness centrality* of two nodes at left putamen, one in the Primary Visual Network and another in the Somatomotor Network (Table S3). *Clustering Coefficient*: A measure of the interconnectivity of neighbors of a node. Age was negatively associated with *clustering coefficient* of nodes across 12 CAB-NP networks, particularly in the Secondary Visual Network, Somatomotor Network, and Cingulo-Opercular Network (Table S5). *Participation Coefficient*: A measure of a node is connected to nodes in other networks. Age was positively associated with *participation coefficient* of nodes in the Auditory Network, Cingulo-Opercular Network, Somatomotor Network, and Secondary Visual Network. On the other hand, age was negatively associated with two nodes in the Primary Visual Network (Table S6). Regarding global graph measures, age was negatively associated with *modularity*, the extent of network nodes forming functional networks (*z* = −2.31, *p_FWE_* = .0096) (Figure 5 (a)) and was positively associated with the *average shortest path length*, the average shortest path length between all possible pairs of nodes, (*z* = 3.27, *p_FWE_* = .0004) (Figure 5 (c)).

**Figure 4.**
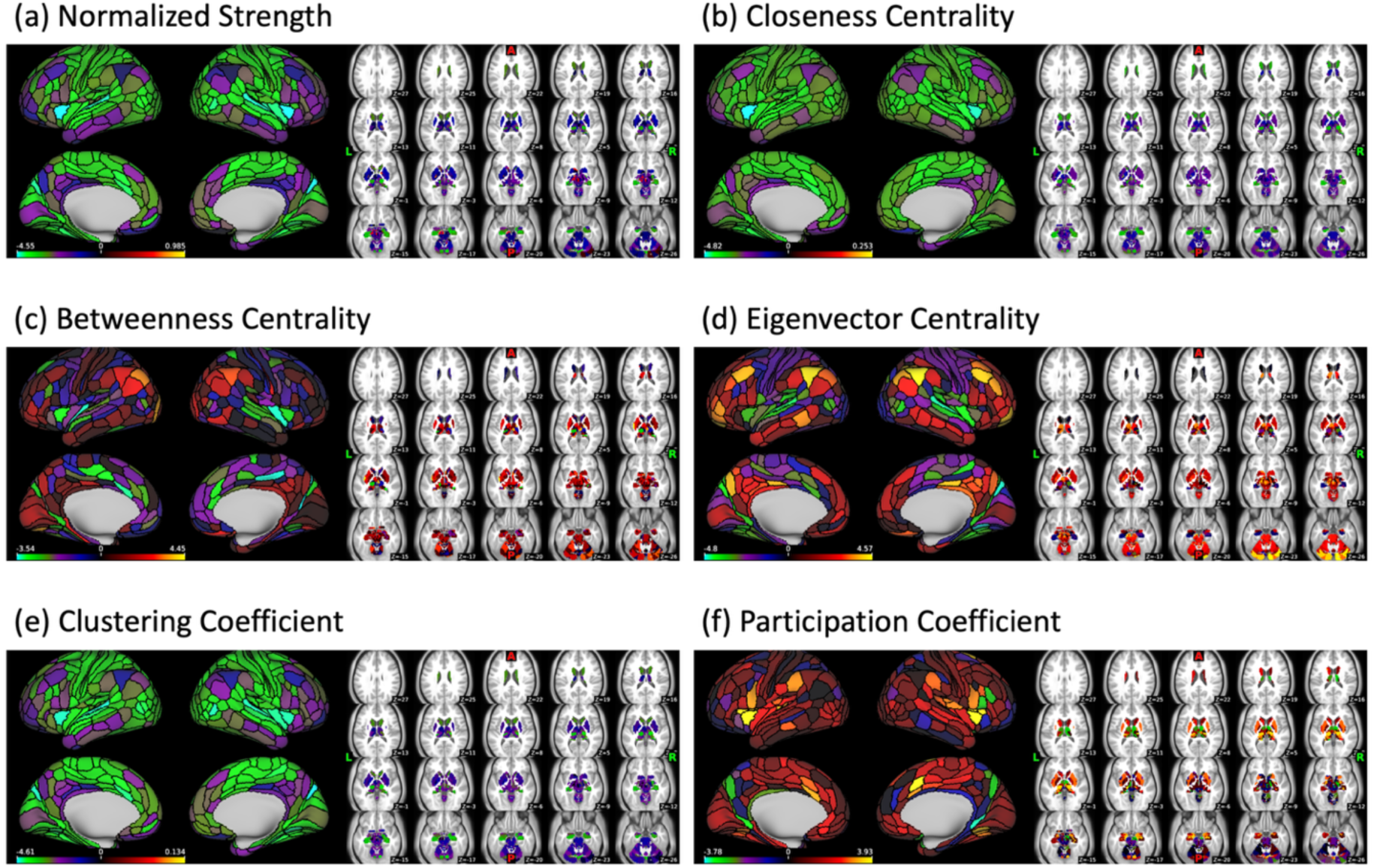
Age and Graph-Based Brain Functional Connectivity Association Maps. Color bar shows the z-score from Aspin-Welch’s test for using exchangeability blocks to adjusted for four scanning sites and variance was estimated for each block, implemented in FSL PALM.

**Figure 5.**
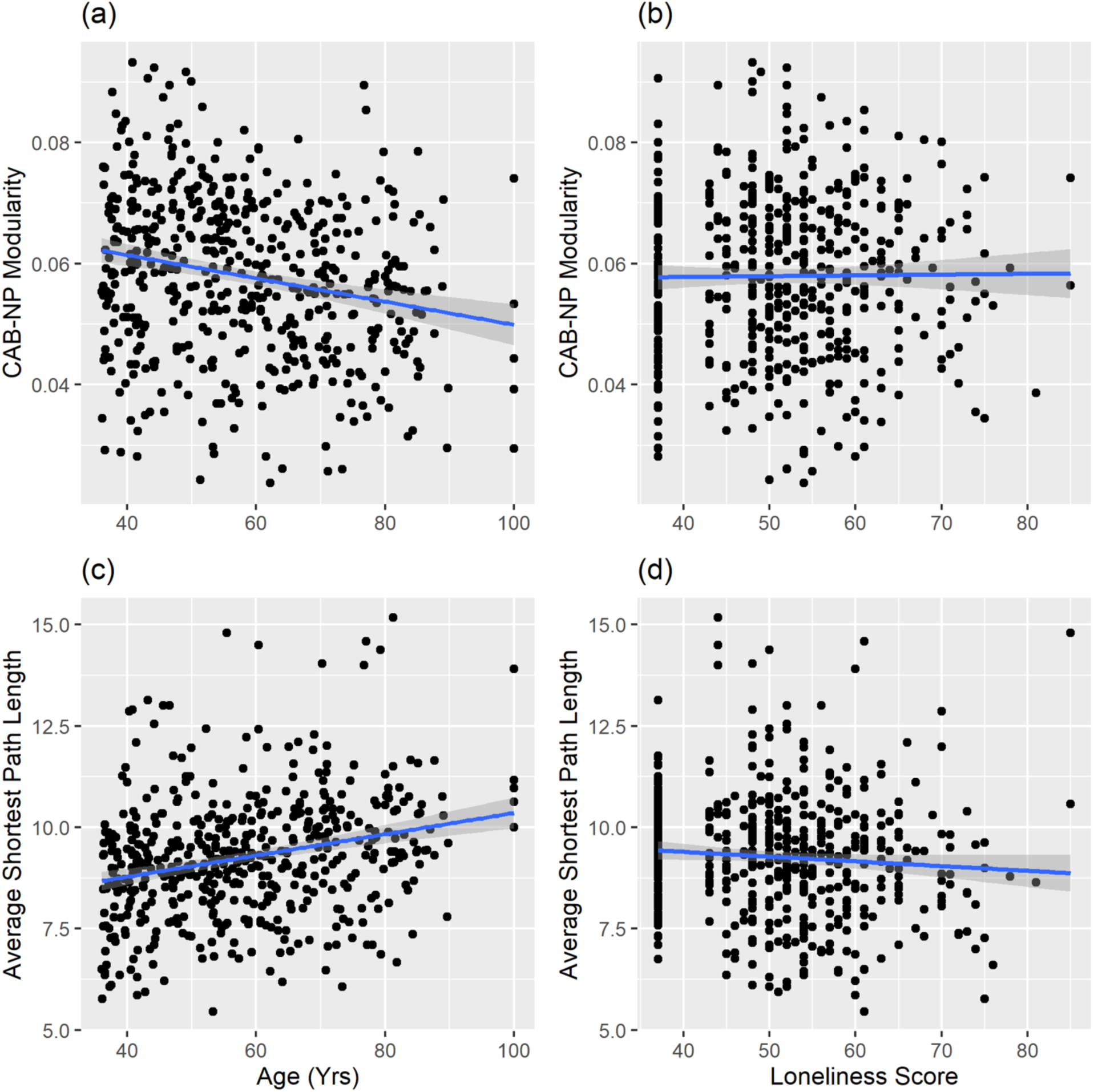
Association of Age and Global Graph-Based Brain Functional Connectivity. Modularity association with (a) age and (b) loneliness. Average shortest path length with (c) age and (d) loneliness.

**Table 3:**
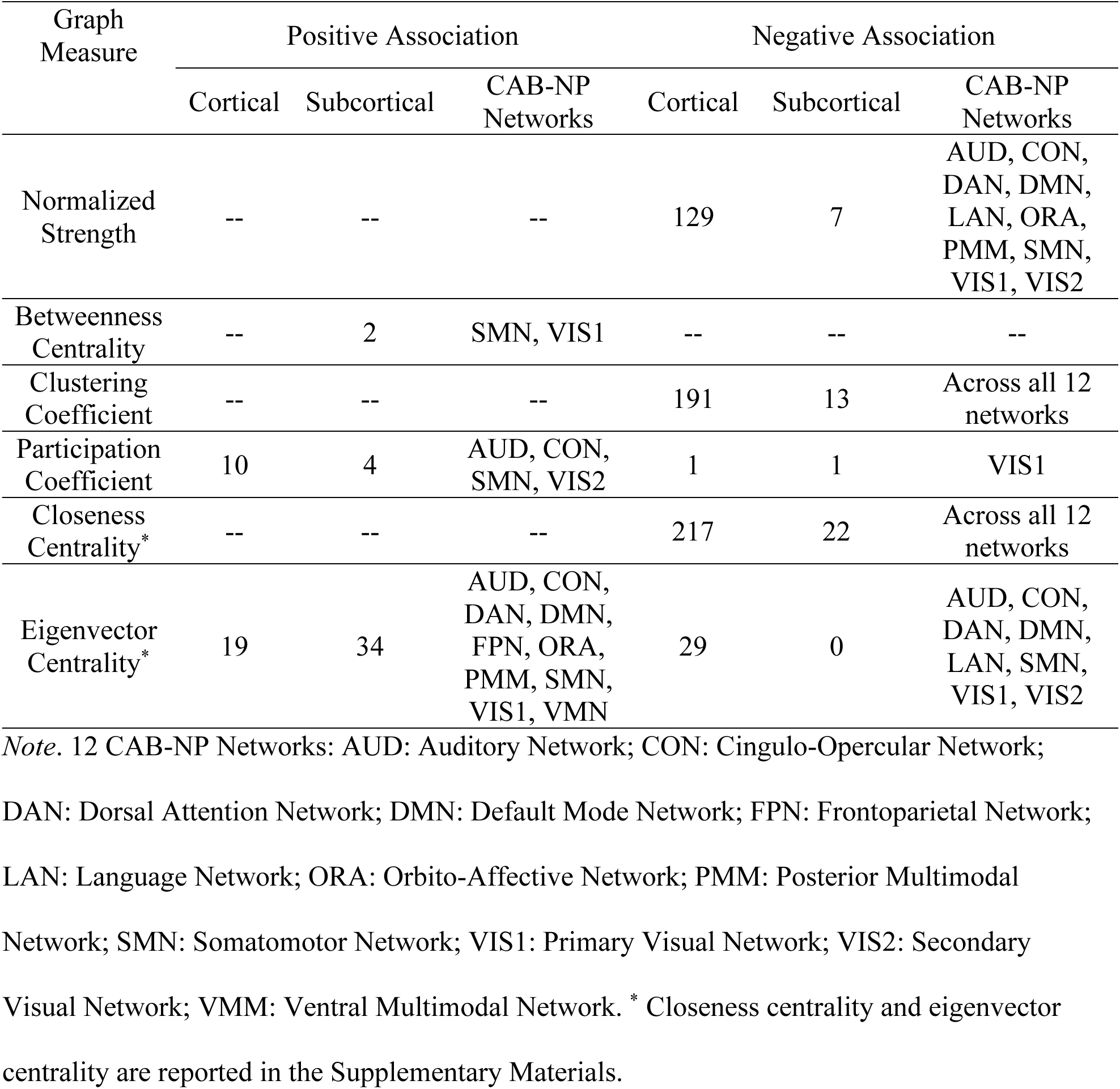
Summary of Age and Graph-Based Brain Functional Connectivity Results at Network-level.

| Graph Measure | Positive Association |  |  | Negative Association |  |  |
| --- | --- | --- | --- | --- | --- | --- |
|  | Cortical | Subcortical | CAB-NP Networks | Cortical | Subcortical | CAB-NP Networks |
| Normalized Strength | -- | -- | -- | 129 | 7 | AUD, CON, DAN, DMN, LAN, ORA, PMM, SMN, VIS1, VIS2 |
| Betweenness Centrality | -- | 2 | SMN, VIS1 | -- | -- | -- |
| Clustering Coefficient | -- | -- | -- | 191 | 13 | Across all 12 networks |
| Participation Coefficient | 10 | 4 | AUD, CON, SMN, VIS2 | 1 | 1 | VIS1 |
| Closeness Centrality* | -- | -- | -- | 217 | 22 | Across all 12 networks |
| Eigenvector Centrality* | 19 | 34 | AUD, CON, DAN, DMN, FPN, ORA, PMM, SMN, VIS1, VMN | 29 | 0 | AUD, CON, DAN, DMN, LAN, SMN, VIS1, VIS2 |

*Note.* 12 CAB-NP Networks: AUD: Auditory Network; CON: Cingulo-Opercular Network;
DAN: Dorsal Attention Network; DMN: Default Mode Network; FPN: Frontoparietal Network;
LAN: Language Network; ORA: Orbito-Affective Network; PMM: Posterior Multimodal
Network; SMN: Somatomotor Network; VIS1: Primary Visual Network; VIS2: Secondary
Visual Network; VMM: Ventral Multimodal Network. \* Closeness centrality and eigenvector
centrality are reported in the Supplementary Materials.

## Discussion

Our study investigated (1) socio-demographic factors associated with loneliness and (2) brain functional connectivity associated with loneliness among individuals aged between 36–100 years old, by leveraging the Lifespan HCP Aging dataset and implemented graph theory.

### Loneliness and Socio-Demographic Associations

Age was negatively associated with loneliness, independent of other socio-demographic factors. Although loneliness is reported to increase with age or follow a U-shaped pattern across the lifespan (Luhmann & Hawkley, 2016; Qualter et al., 2015), a negative age-loneliness association has been reported in other studies using similar middle-to-older adult samples (Spreng et al., 2020; C. Wilson & Moulton, 2010). It is important to note that the HCP-Aging cohort excluded individuals with cognitive impairment and screened for cognitive ability. This selection process may result in recruiting healthier, more socially resilient older adults, which may attenuate or even reverse the positive age-loneliness association typically observed in more representative populations. Findings regarding the age-loneliness relationship should therefore be interpreted with caution.

The age-related difference in loneliness may reflect distinct social needs and social norms at different age stages (Luhmann & Hawkley, 2016; Qualter et al., 2015). Considering multiple socio-demographic factors, the results showed that being of younger age, male, unmarried, and living alone was associated with higher levels of loneliness, consistent with earlier work (Beutel et al., 2017; Distel et al., 2010; Dykstra et al., 2005; Ge et al., 2017; Hawkley et al., 2005). No significant association was found with employment status or education. Sex differences emerged only after adjusting for other factors, whereas the association with income weakened. These observations emphasize the importance of including diverse socio-demographic factors and exploring their interactions in future loneliness research.

### Loneliness and Graph-Based Brain Functional Connectivity

We found no significant association between loneliness and any functional connectivity measures, nor any significant interaction between loneliness and age. To further probe whether age-related changes in global network properties (modularity, average shortest path length), Supplement Figure S5 illustrates participants in the highest and lowest loneliness tertiles, showing broadly similar slopes across groups. When accounting for socio-demographic factors that were significantly associated with loneliness (i.e., sex, marital status, and household status), the results showed that the interaction between loneliness and age was associated with the *clustering coefficient* of left caudate in the Default Mode Network and right cerebellum in the Frontoparietal Network. For both nodes, the association with the *clustering coefficient* is positive for younger individuals (∼ under 65 years old), but negative for older individuals aged (∼ over 65 years old). This interaction reflects an age-dependent influence on the association between loneliness experience and brain network organization. Thus, younger individuals show increased local interconnectivity, but older individuals show decreased local interconnectivity associated with loneliness. The involvement of DMN and FPN suggest that loneliness might have impact on neural systems that involve both internal mental process and executive control process (Andrews-Hanna et al., 2014; Dosenbach et al., 2007), and its impact differs across different ages. Furthermore, the interplay of age and loneliness on brain functional organization emerges while taking account for socio-demographic factors including sex, marital status, and household status, which emphasizes the need to consider important socio-demographic factors of loneliness in future investigations.

Contrary to expectations informed by prior resting-state fMRI studies, no significant main effect of loneliness was observed. Most prior studies employed seed-based or pairwise functional connectivity approaches (Brilliant T et al., 2022; Feng et al., 2019; Spreng et al., 2020). We are aware of only one other study that measured graph modularity (Mwilambwe-Tshilobo et al., 2019), and which reported a negative association. Differences from the present findings may reflect several methodological factors: first, the two studies differed in sample characteristics, with the current study including a broader age range (36-100 years), whereas Mwilambwe-Tshilobo et al. focused on a younger cohort (23-37 years). Second, the studies used different network parcellation schemes, with the present work employing 12-network CAB-NP template including cortical and subcortical regions (Ji et al., 2019), whereas the prior study used a seven-network cortical-only parcellation (Yeo et al., 2011). Finally, differences in analytic focus may also contribute, with prior studies primarily quantifying pairwise connectivity between regions or networks, whereas graph-theoretical measures capture higher-order properties of whole-brain network organization derived from these connectivity patterns.

In addition, the present analyses used stringent family-wise error correction across nodes within each graph metric, which may reduce sensitivity to detect subtle effects while prioritizing control of false positives in high-dimensional data. This approach was chosen to minimize the risk of spurious findings given the large number of nodal comparisons. Future work combining graph-theoretic approahces with targeted seed-based or network-level connectivity analyses may help clarify whether previously reported connectivity effects (e.g., within default network or between-network altrerations) are also reflected in global network topology. Overall, the present study may complement prior pairwise connectivity work by extending the investigation of loneliness-related neural differences to higher-order properties of whole-brain network organization.

### Age and Graph-Based Brain Functional Connectivity

In contrast to the null results observed with loneliness, age showed significant associations with all functional connectivity measures. Age was positively associated with the *average shortest path length*, indicating that the averaged shortest path between connectivity pairs lengthens with age. This connectivity pattern may represent lower global efficiency and a higher cost for information integration among older people (Achard & Bullmore, 2007; Sala-Llonch et al., 2014). At the nodal level, age was associated with lower *connectivity strength* and *closeness centrality* of nodes across the whole brain, indicating a lower degree of connectivity and directed connections among older people. These findings suggest that there could be a decrease in the overall functional connectivity with age, particularly in networks related to sensory-motor, task maintenance, and self-generated thought processes (Andrews-Hanna et al., 2014; Dosenbach et al., 2007). Additionally, age was negatively associated with *modularity*, indicating lower modular organization among older people (Betzel et al., 2014; Cao et al., 2014; Geerligs et al., 2015). This finding, to some extent, corresponds to the negative association between age and *clustering coefficient*, indicating lower ability to form highly interconnected and functionally specialized brain clusters. Conversely, age was positively associated with *participation coefficient*, indicating that these nodes among older people connect more with nodes in other networks, rather than within the same network. These results suggest a less segregated brain network organization among older people (Chan et al., 2014; Geerligs et al., 2015; Stumme et al., 2020). Such a connectivity pattern has previously been associated with less efficient information processing and with poorer cognitive functions, such as non-verbal memory and attention (Chan et al., 2014; Stumme et al., 2020).

These age-associated differences were not homogeneous across the whole brain. Besides its positive association with *participation coefficient*, age was negatively associated with *participation coefficient* of two nodes in the Primary Visual Network, indicating these nodes have lower connectivity with other networks among older people. Heterogeneous associations were also observed with *eigenvector centrality*. Age was positively associated with *eigenvector centrality* of several nodes, particularly in the Frontoparietal Network, suggesting increased influence of these nodes within the Frontoparietal Network among older people. Age was also positively associated with *betweenness centrality* of two left putamen nodes in the Primary Visual Network and the Somatomotor Network, indicating more information passes through these two nodes among older people. The increased influence of these nodes may reflect a compensatory mechanism for age-associated changes (Reuter-Lorenz & Cappell, 2008). Conversely, age was negatively associated with *eigenvector centrality* of nodes particularly in the Auditory and Secondary Visual networks, indicating decreased influence. The weaker influence of these nodes may indicate disrupted functional integration between sensory and other brain networks with aging. The mixed age-related positive and negative associations reveal heterogeneous alterations in brain network functional connectivity. Future research is needed to delineate these variations in age-associated brain functional connectivity and their links to cognitive and behavioral abilities, determining whether they serve as compensatory mechanisms or disruptive impairments.

### Limitations

It is important to acknowledge certain limitations while interpreting the results of this study. First, our study was based on cross-sectional data, which limits the ability to establish causal relationships. Second, loneliness was measured using the NIH Toolbox Loneliness Survey, which captures the *Isolation* dimension of loneliness (Cyranowski et al., 2013), but not the other two dimensions of *Relational* and of *Collective Connectedness* (Hawkley et al., 2005). Third, only short-term (past month) experience of loneliness was captured in the Loneliness Survey, whereas prior work reported that persistent, but not transient, loneliness was associated with measures of cognitive performance and neural measures (Tao et al., 2022). These measurement characteristics may be particularly consequential in a sample spanning a wide age range such as the HCP-Aging cohort. Younger adults may report higher levels of transient situational loneliness related to life transitions (e.g., career relocation), which may not reflect stable patterns of social disconnection. Conversely, older adults may report lower current loneliness despite reduced social networks due to long-term adaptation to changes in social relationships. Fourth, depressive symptoms were not included as a covariate. Given that loneliness and depression are highly correlated (Chen et al., 2023), and depression has documented effects on brain functional connectivity (Zhang et al., 2025). Future work incorporating affective symptoms may help clarify the specificity of loneliness-related associations with brain network organization. Fifth, whether the negative connectivity weight carries meaningful neurobiological information is an ongoing discussion (Fox et al., 2009; Murphy et al., 2009). Due to the ambiguity of negative weights interpretation, our study took the absolute value of connectivity weights. However, it is important to acknowledge that positive and negative connectivity could carry different neurobiological meanings and could potentially impose confounds for graph measures (G. Chen et al., 2011; Rubinov & Sporns, 2011; Schwarz & McGonigle, 2011). For exploratory purpose, our study conducted separate analyses to evaluate positive-only and negative-only weights, respectively. Results from the positive-only measures were mostly consistent with the absolutized results, whereas the negative-only results were greatly attenuated (Supplementary Materials). Future research should carefully assess and implement alternative approaches, such as binarization with careful consideration of the choice of threshold (Schwarz & McGonigle, 2011) and evaluate positive and negative weights separately (Geerligs et al., 2015; Rubinov & Sporns, 2011). Sixth, our study used total gray matter volume to account for age-related brain structural differences. However, studies have shown a close relationship between functional connectivity and structural connectivity (Deco et al., 2010; Hermundstad et al., 2013; Honey et al., 2009). Betzel et al. (2014) observed that age-related functional connectivity could be accounted by structural connectivity, particularly those short-range structurally connected paths. Finally, drawing meaningful inferences from resting-state data becomes challenging without incorporating behavioral-level functional assessments. This could be particularly challenging for a subjective and dimensionally complex construct like loneliness. Given the well-established association between loneliness and a range of adaptive and maladaptive cognitive and behavioral functions, it is plausible that these processes could play a role as moderators or mediators in the relationship between loneliness and neural measures. Moreover, utilizing methods to manipulate experienced loneliness (e.g., by temporary social isolation) could offer a powerful approach for direct functional inference of the association between loneliness and brain activity. Taken together, future research should incorporate longitudinal, multi-method, and cross-modality approaches to address multi-dimensional nature and temporal aspect of loneliness to gain a more comprehensive understanding of this complex psychological construct.

### Strengths

Our study investigated the association between loneliness and age across a broad age range from thirty-six to one-hundred years old, expanding on prior research that focused mainly on younger adults or on older adults, respectively. Most prior neuroimaging studies of loneliness only included a few socio-demographic factors and usually treated these as variables of no interest. By accounting for loneliness-associated socio-demographic factors in both behavioral and brain functional connectivity analyses, our study underscores the importance of considering these factors in research on loneliness. Our study expanded upon prior studies that used cortical mapped parcellations by utilizing CAB-NP network parcellation (Ji et al., 2019) to investigate brain functional connectivity across cortical and subcortical structures. In addition, we are aware of only one RSFC study on loneliness that had utilized a graph approach. Incorporating resting-state functional imaging and graph theory offers a powerful tool for examining both the local and global organization of brain networks. By examining multiple graph measures, our study offers a comprehensive analysis of brain network organization associated with loneliness.

### Conclusion

Our study observed (1) higher levels of loneliness were associated with younger age, being male, unmarried status, and living alone, (2) loneliness did not show any association with local or global graph measures, (3) interaction between loneliness and age was associated with local interconnectivity in DMN and FPN, while accounting for important socio-demographic factors, (4) age was substantially associated with graph measures, with older age linked to lower functional connectivity, reduced global efficiency, and diminished brain network modularity, (5) age also showed varying associations with graph measures in different brain networks. In conclusion, our study reveals weak neural correlates of loneliness. Although unexpected, it underscores the complexity of the subjective and multi-dimensional nature of loneliness. It highlights the limitations of current methods and prompts that future research should incorporate longitudinal designs, diverse loneliness measures beyond self-report, and multi-modal neuroimaging techniques.

## Funding & Acknowledgements

This work received no specific grant from any funding agency. We would like to thank the first author’s dissertation committee, Dr. Hoi-Chung Leung, Dr. Brady Nelson, and Dr. Roman Kotov, for their invaluable guidance and support to shape the current work. We would also like to thank the undergraduate research assistants, Richard Adam, Ryan George, Nyomi Sutherland, Farhan Noor, Victoria Bayevskiy, and Sisira Gajjala, for their contributions on conducting quality assessment of the neuroimaging data.

## Data Availability

The data that support the findings of this study are openly available in the Lifespan Human Connectome Project. Data collection and sharing for the Lifespan Human Connectome Project Aging was supported by the National Institute On Aging of the National Institutes of Health under Award Number U01AG052564 and by funds provided by the McDonnell Center for Systems Neuroscience at Washington University in St. Louis. The HCP-Aging 2.0 Release data used in this report came from DOI: <u>10.15154/1520707</u>. Data and/or research tools used in the preparation of this manuscript were obtained from the National Institute of Mental Health (NIMH) Data Archive (NDA). NDA is a collaborative informatics system created by the National Institutes of Health to provide a national resource to support and accelerate research in mental health. Dataset identifier(s): 10.15154/g2tv-f889. This manuscript reflects the views of the authors and may not reflect the opinions or views of the NIH or of the Submitters submitting original data to NDA.

## Author Contribution Statement

**Yen-Wen Chen:** Conceptualization, Data curation, Formal analysis, Investigation, Methodology, Project Administration, Resources, Validation, Visualization, Writing - original draft, Writing - review & editing.

**Turhan Canli:** Conceptualization, Investigation, Methodology, Project Administration, Resources, Supervision, Writing - review & editing.

