## Supplementary Materials for "Social Disconnection in the Brain: Loneliness and Age across Networks using Graph Theory"

### Supplementary Graph-Theoretical Measures and Results

Primary analyses focused on representative graph metrics: degree centrality, betweenness centrality, shortest path length, clustering coefficient, and modularity. Additional related metrics (closeness centrality and eigenvector centrality) are reported here to reduce redundancy among highly correlated graph measures.

#### *Closeness Centrality*

*Closeness centrality* quantifies a node's proximity to all other nodes in the network, calculated as the reciprocal of the average shortest path length to a node over all other connected nodes. High *closeness centrality* indicates topologically central or influential nodes within the network (Fornito et al., 2016). No significant main effect of loneliness, nor an interaction of loneliness and age, was revealed with this metric. On the other hand, age was negatively associated with *closeness centrality* of nodes across all 12 CAB-NP networks, particularly in the Secondary Visual Network, Cingulo-Opercular Network, Somatomotor Network, and Default Mode Network (Table S2).

#### *Eigenvector Centrality*

*Eigenvector centrality* quantifies a node's importance by accounting for both the quantity and quality of its connections, defined as the node's entry in the eigenvector corresponding to the largest eigenvalue of the matrix. Higher values indicate a node's connection to other well-connected nodes, highlighting their central roles in the network (Fornito et al., 2016; Van Duinkerken et al., 2017). No significant main effect of loneliness, nor an interaction of loneliness and age, was revealed with this metric. On the other hand, age was positively associated with *eigenvector centrality* of nodes across 10 CAB-NP networks, particularly in the Frontoparietal

Network. On the other hand, age was negatively associated with nodes across eight CBA-NP networks, particularly in the Auditory Network and Secondary Visual Network (Table S4).

**Supplementary Analyses of Positive-Only and Negative-Only Functional Connectivity**

To obtain positive-only connectivity weights, the negative weighted edges were omitted, and for negative-only connectivity weights, the positive weighted edges were omitted (Rubinov & Sporns, 2011). For both positive-only and negative-only graph-based measures, no association with loneliness, nor the interaction between loneliness and age was observed. Regarding positive-only graph-based measures (Table S7, Figure S1), the association between age and nodal graph-based measures were consistent with absolutized measures. Age also showed consistent positive association with the *average shortest path length* ($z$ = 3.53, $p_{FWE}$ = .0002). However, the association between age and *modularity* was not significant ($z$ = -.65, $p_{FWE}$ = .5052) (Figure S2). Regarding negative-only graph-based measures (Table S8, Figure S3), the associations between age and nodal graph-based measures were significantly attenuated. On the other hand, age was positively associated with the *average shortest path length* ($z$ = 2.57, $p_{FWE}$ = .0060), whereas no association with *modularity* ($z$ = 1.64, $p_{FWE}$ = .0822) (Figure S4).

### List of Supplementary Tables

Supplementary Table S1 Age and Nodal Strength Association Results

Supplementary Table S2 Age and Nodal Closeness Centrality Association Results

Supplementary Table S3 Age and Nodal Betweenness Centrality Association Results

Supplementary Table S4 Age and Nodal Eigenvector Centrality Association Results

Supplementary Table S5 Age and Nodal Clustering Coefficient Association Results

Supplementary Table S6 Age and Nodal Participation Coefficient Association Results

Supplementary Table S7 Summary of Age and Graph-Based Brain Functional Connectivity Results: Positive-Only Connectivity Weights

Supplementary Table S8 Summary of Age and Graph-Based Brain Functional Connectivity Results: Negative-Only Connectivity Weights

### List of Supplementary Figures

Supplementary Figure S1 Age and Graph-Based Brain Functional Connectivity Association Maps: Positive-Only Connectivity Weights. Color bar shows the z-score from Aspin-Welch's test for using exchangeability blocks to adjusted for four scanning sites and variance was estimated for each block, implemented in FSL PALM

Supplementary Figure S2 Association of Age and Global Graph-Based Brain Functional Connectivity: Positive-Only Connectivity Weights. Modularity association with (a) age and (b) loneliness. Average shortest path length with (c) age and (d) loneliness

Supplementary Figure S3 Age and Graph-Based Brain Functional Connectivity Association Maps: Negative-Only Connectivity Weights. Color bar shows the z-score from Aspin-Welch's test for using exchangeability blocks to adjusted for four scanning sites and variance was estimated for each block, implemented in FSL PALM

Supplementary Figure S4 Association of Age and Global Graph-Based Brain Functional Connectivity: Negative-Only Connectivity Weights. Modularity association with (a) age and (b) loneliness. Average shortest path length with (c) age and (d) loneliness

Supplementary Figure S5 Figure is shown for illustrative purposes, depicting age-related trajectories in the loneliest (top tertile) and most socially connected (bottom tertile) individuals. The formal test of the loneliness × age interaction was conducted in PALM across the full sample (N = 512) using loneliness as a continuous variable.

**Supplementary Table S1**

***Age and Nodal Strength Association Results***

| CAB-NP Label[a] | Glasser Label[b] | z[c] | -log10($p_{FWE}$) |
|---|---|---|---|
| Cingulo-Opercular-47_L-Ctx | Middle Insular Area | -4.55 | 3.7 |
| Cingulo-Opercular-20_R-Ctx | Middle Insular Area | -4.5 | 3.7 |
| Visual2-02_R-Ctx | Sixth Visual Area | -4.49 | 3.7 |
| Auditory-07_R-Ctx | Auditory 4 Complex | -4.39 | 3.7 |
| Default-33_R-Ctx | ParaHippocampal Area 2 | -4.39 | 3.7 |
| Auditory-15_L-Ctx | Auditory 4 Complex | -4.36 | 3.7 |
| Cingulo-Opercular-21_R-Ctx | Frontal OPercular Area 1 | -4.34 | 3.7 |
| Cingulo-Opercular-25_R-Ctx | Area Posterior Insular 1 | -4.29 | 3.7 |
| Visual2-29_L-Ctx | Sixth Visual Area | -4.28 | 3.7 |
| Cingulo-Opercular-22_R-Ctx | Frontal OPercular Area 3 | -4.24 | 3.7 |
| Auditory-04_R-Ctx | ParaBelt Complex | -4.23 | 3.7 |
| Cingulo-Opercular-18_R-Ctx | Posterior Insular Area 2 | -4.21 | 3.7 |
| Visual2-01_R-Ctx | Medial Superior Temporal Area | -4.21 | 3.7 |
| Auditory-12_L-Ctx | ParaBelt Complex | -4.16 | 3.7 |
| Visual2-20_R-Ctx | VentroMedial Visual Area 1 | -4.16 | 3.7 |
| Cingulo-Opercular-38_L-Ctx | Anterior 24 prime | -4.14 | 3.7 |
| Dorsal-Attention-19_L-Ctx | ParaHippocampal Area 3 | -4.13 | 3.7 |
| Visual2-53_L-Ctx | VentroMedial Visual Area 2 | -4.13 | 3.7 |
| Dorsal-Attention-07_R-Ctx | ParaHippocampal Area 3 | -4.11 | 3.7 |
| Language-06_R-Ctx | Auditory 5 Complex | -4.09 | 3.7 |
| Cingulo-Opercular-44_L-Ctx | Area PFcm | -4.07 | 3.4 |
| Visual1-06_L-Ctx | Dorsal Transitional Visual Area | -4.05 | 3.4 |
| Cingulo-Opercular-39_L-Ctx | Area p32 prime | -4.05 | 3.4 |
| Visual2-46_L-Ctx | Area V6A | -4.05 | 3.4 |
| Visual1-03_R-Ctx | Dorsal Transitional Visual Area | -4.04 | 3.4 |
| Cingulo-Opercular-01_R-Ctx | Frontal Eye Fields | -4.03 | 3.4 |
| Visual2-47_L-Ctx | VentroMedial Visual Area 1 | -4.02 | 3.4 |
| Cingulo-Opercular-04_R-Ctx | Area 5m ventral | -4.02 | 3.4 |

| | | | |
|---|---|---|---|
| Visual2-08_R-Ctx | Seventh Visual Area | -4.01 | 3.4 |
| Cingulo-Opercular-52_L-Ctx | Area Posterior Insular 1 | -4.01 | 3.4 |
| Visual2-28_L-Ctx | Medial Superior Temporal Area | -4.01 | 3.4 |
| Language-19_L-Ctx | Auditory 5 Complex | -3.99 | 3.4 |
| Visual2-26_R-Ctx | VentroMedial Visual Area 2 | -3.99 | 3.4 |
| Default-25_R-Ctx | ParaHippocampal Area 1 | -3.98 | 3.4 |
| Language-07_R-Ctx | Area STSd posterior | -3.96 | 3.4 |
| Cingulo-Opercular-16_R-Ctx | Area 43 | -3.93 | 3.4 |
| Default-64_L-Ctx | ParaHippocampal Area 1 | -3.92 | 3.4 |
| Default-71_L-Ctx | ParaHippocampal Area 2 | -3.91 | 3.4 |
| Cingulo-Opercular-17_R-Ctx | Area PFcm | -3.91 | 3.4 |
| Auditory-03_R-Ctx | Area TA2 | -3.91 | 3.4 |
| Visual2-16_R-Ctx | Area Lateral IntraParietal ventral | -3.91 | 3.4 |
| Visual2-27_R-Ctx | Ventral Visual Complex | -3.91 | 3.4 |
| Cingulo-Opercular-49_L-Ctx | Frontal OPercular Area 3 | -3.9 | 3.4 |
| Visual2-19_R-Ctx | Area V6A | -3.89 | 3.22 |
| Cingulo-Opercular-48_L-Ctx | Frontal OPercular Area 1 | -3.87 | 3.1 |
| Visual2-42_L-Ctx | Middle Temporal Area | -3.87 | 3.1 |
| Cingulo-Opercular-43_L-Ctx | Area 43 | -3.86 | 3.1 |
| Visual2-38_L-Ctx | Area V3B | -3.84 | 3.1 |
| Visual2-15_R-Cerebellum | | -3.84 | 3.1 |
| Visual2-22_R-Ctx | Area V4t | -3.82 | 3 |
| Auditory-06_R-Ctx | Lateral Belt Complex | -3.82 | 3 |
| Cingulo-Opercular-30_L-Ctx | Frontal Eye Fields | -3.81 | 3 |
| Cingulo-Opercular-06_R-Ctx | Supplementary and Cingulate Eye Field | -3.8 | 3 |
| Language-20_L-Ctx | Area STSd anterior | -3.79 | 3 |
| Visual2-34_L-Ctx | Area V3A | -3.78 | 2.74 |
| Visual2-54_L-Ctx | Ventral Visual Complex | -3.77 | 2.74 |
| Posterior-Multimodal-02_R-Ctx | Superior Temporal Visual Area | -3.77 | 2.74 |
| Somatomotor-08_R-Ctx | Area 7PC | -3.77 | 2.74 |
| Somatomotor-13_R-Ctx | Area 6mp | -3.77 | 2.74 |

| | | | |
|---|---|---|---|
| Default-63_L-Ctx | Hippocampus | -3.76 | 2.7 |
| Visual2-43_L-Ctx | Area Lateral IntraParietal ventral | -3.76 | 2.59 |
| Auditory-14_L-Ctx | Lateral Belt Complex | -3.76 | 2.59 |
| Cingulo-Opercular-31_L-Ctx | Area 5m ventral | -3.75 | 2.59 |
| Dorsal-Attention-04_R-Ctx | Area 6 anterior | -3.75 | 2.59 |
| Visual2-15_R-Ctx | Middle Temporal Area | -3.75 | 2.59 |
| Cingulo-Opercular-03_R-Ctx | PeriSylvian Language Area | -3.74 | 2.59 |
| Somatomotor-15_R-Ctx | Area OP4/PV | -3.72 | 2.52 |
| Visual2-25_R-Ctx | Area Lateral Occipital 3 | -3.72 | 2.52 |
| Cingulo-Opercular-27_R-Ctx | Para-Insular Area | -3.72 | 2.52 |
| Default-18_R-Ctx | Area 9 anterior | -3.72 | 2.52 |
| Language-13_L-Ctx | Superior Temporal Visual Area | -3.72 | 2.52 |
| Visual2-35_L-Ctx | Seventh Visual Area | -3.71 | 2.47 |
| Visual2-07_R-Ctx | Area V3A | -3.7 | 2.44 |
| Cingulo-Opercular-32_L-Ctx | Area 23c | -3.7 | 2.44 |
| Visual2-21_R-Ctx | VentroMedial Visual Area 3 | -3.69 | 2.44 |
| Visual2-09_R-Ctx | IntraParietal Sulcus Area 1 | -3.68 | 2.44 |
| Language-01_R-Ctx | Area 55b | -3.68 | 2.44 |
| Somatomotor-18_R-Ctx | RetroInsular Cortex | -3.67 | 2.36 |
| Language-08_R-Ctx | Area TemporoParietoOcci pital Junction 1 | -3.67 | 2.32 |
| Somatomotor-19_R-Ctx | Frontal OPercular Area 2 | -3.67 | 2.3 |
| Somatomotor-19_R-Hippocampus | | -3.67 | 2.3 |
| Visual2-48_L-Ctx | VentroMedial Visual Area 3 | -3.66 | 2.3 |
| Posterior-Multimodal-03_R-Ctx | Area TemporoParietoOcci pital Junction 2 | -3.66 | 2.28 |
| Language-16_L-Ctx | Area IFJa | -3.65 | 2.27 |
| Orbito-Affective-01_R-Ctx | Pirform Cortex | -3.65 | 2.27 |
| Visual2-30_L-Ctx | Second Visual Area | -3.64 | 2.27 |
| Dorsal-Attention-09_L-Cerebellum | | -3.64 | 2.27 |
| Dorsal-Attention-20_L-Ctx | Area TE2 posterior | -3.64 | 2.27 |
| Dorsal-Attention-14_L-Ctx | Medial IntraParietal Area | -3.64 | 2.27 |

| | | | |
|---|---|---|---|
| Somatomotor-28_R-Thalamus | | -3.64 | 2.27 |
| Posterior-Multimodal-07_L-Ctx | Area TemporoParietoOcci pital Junction 3 | -3.64 | 2.25 |
| Visual2-36_L-Ctx | IntraParietal Sulcus Area 1 | -3.63 | 2.24 |
| Posterior-Multimodal-10_L-Cerebellum | | -3.62 | 2.19 |
| Cingulo-Opercular-11_R-Ctx | Area p32 prime | -3.61 | 2.15 |
| Visual1-02_R-Ctx | ProStriate Area | -3.61 | 2.15 |
| Visual2-08_L-Cerebellum | | -3.6 | 2.13 |
| Auditory-11_L-Ctx | Area TA2 | -3.59 | 2.11 |
| Default-15_R-Ctx | Area 9 Posterior | -3.59 | 2.11 |
| Visual1-05_L-Ctx | ProStriate Area | -3.59 | 2.1 |
| Cingulo-Opercular-45_L-Ctx | Posterior Insular Area 2 | -3.59 | 2.09 |
| Default-37_R-Ctx | Area STSv anterior | -3.59 | 2.09 |
| Default-53_L-Ctx | Area 9 Posterior | -3.59 | 2.09 |
| Visual2-06_R-Ctx | Eighth Visual Area | -3.58 | 2.05 |
| Visual2-17_R-Ctx | Ventral IntraParietal Complex | -3.58 | 2.03 |
| Auditory-10_L-Ctx | RetroInsular Cortex | -3.56 | 2.01 |
| Somatomotor-36_L-Ctx | Area OP1/SII | -3.56 | 2 |
| Somatomotor-28_L-Ctx | Area 7PC | -3.56 | 1.98 |
| Visual2-03_R-Ctx | Second Visual Area | -3.56 | 1.98 |
| Language-02_R-Ctx | Superior Frontal Language Area | -3.55 | 1.98 |
| Cingulo-Opercular-10_R-Ctx | Anterior 24 prime | -3.55 | 1.97 |
| Default-26_R-Ctx | Area STSd anterior | -3.55 | 1.97 |
| Visual2-37_L-Ctx | Fusiform Face Complex | -3.54 | 1.96 |
| Visual2-31_L-Ctx | Third Visual Area | -3.54 | 1.96 |
| Dorsal-Attention-23_L-Ctx | Area IntraParietal 0 | -3.54 | 1.96 |
| Posterior-Multimodal-04_R-Ctx | Area TemporoParietoOcci pital Junction 3 | -3.54 | 1.94 |
| Cingulo-Opercular-07_R-Ctx | Area 6m anterior | -3.54 | 1.94 |
| Language-17_L-Ctx | Area IFSp | -3.52 | 1.89 |
| Visual2-10_R-Ctx | Fusiform Face Complex | -3.52 | 1.89 |
| Default-14_R-Ctx | Area 8B Lateral | -3.51 | 1.87 |
| Somatomotor-33_L-Ctx | Area 6mp | -3.5 | 1.84 |
| Somatomotor-09_R-Ctx | Area 1 | -3.5 | 1.82 |
| Somatomotor-06_R-Ctx | Ventral Area 24d | -3.49 | 1.81 |
| Somatomotor-26_L-Ctx | Ventral Area 24d | -3.49 | 1.81 |
| Cingulo-Opercular-05_R-Ctx | Area 23c | -3.49 | 1.79 |

| | | | |
|---|---|---|---|
| Language-10_L-Ctx | Area 55b | -3.48 | 1.77 |
| Default-47_L-Ctx | Area 10r | -3.48 | 1.76 |
| Visual2-49_L-Ctx | Area V4t | -3.47 | 1.75 |
| Default-76_L-Ctx | Area STSv anterior | -3.46 | 1.73 |
| Cingulo-Opercular-23_R-Ctx | Area PF opercular | -3.45 | 1.69 |
| Somatomotor-12_R-Ctx | Dorsal area 6 | -3.45 | 1.69 |
| Language-21_L-Ctx | Area STSd posterior | -3.45 | 1.69 |
| Visual2-52_L-Ctx | Area Lateral Occipital 3 | -3.44 | 1.69 |
| Somatomotor-10_R-Ctx | Area 2 | -3.44 | 1.67 |
| Language-04_R-Ctx | Area IFJa | -3.43 | 1.67 |
| Somatomotor-23_L-Ctx | Area 5m | -3.43 | 1.63 |
| Cingulo-Opercular-17_R-Cerebellum | | -3.42 | 1.61 |

*Note*. [a]CAB-NP label per Cole-Anticevic Brain Network Parcellation v1.1.6.

[b]Glasser cortical label names were from Glasser, et al. (2016).

[c]*z*-score from Aspin-Welch's test for using exchangeability blocks to adjusted for four scanning sites and variance was estimated for each block, implemented in FSL PALM.

**Supplementary Table S2**

***Age and Nodal Closeness Centrality Association Results***

| CAB-NP Label[a] | Glasser Label[b] | z[c] | -log10($p_{FWE}$) |
|---|---|---|---|
| Cingulo-Opercular-47_L-Ctx | Middle Insular Area | -4.82 | 3.7 |
| Default-33_R-Ctx | ParaHippocampal Area 2 | -4.67 | 3.7 |
| Cingulo-Opercular-20_R-Ctx | Middle Insular Area | -4.62 | 3.7 |
| Cingulo-Opercular-22_R-Ctx | Frontal OPercular Area 3 | -4.49 | 3.7 |
| Cingulo-Opercular-21_R-Ctx | Frontal OPercular Area 1 | -4.45 | 3.7 |
| Dorsal-Attention-07_R-Ctx | ParaHippocampal Area 3 | -4.34 | 3.7 |
| Visual2-53_L-Ctx | VentroMedial Visual Area 2 | -4.33 | 3.7 |
| Cingulo-Opercular-38_L-Ctx | Anterior 24 prime | -4.31 | 3.7 |
| Dorsal-Attention-19_L-Ctx | ParaHippocampal Area 3 | -4.31 | 3.7 |
| Cingulo-Opercular-25_R-Ctx | Area Posterior Insular 1 | -4.31 | 3.7 |
| Cingulo-Opercular-49_L-Ctx | Frontal OPercular Area 3 | -4.29 | 3.7 |
| Cingulo-Opercular-18_R-Ctx | Posterior Insular Area 2 | -4.28 | 3.7 |
| Auditory-15_L-Ctx | Auditory 4 Complex | -4.22 | 3.7 |
| Auditory-04_R-Ctx | ParaBelt Complex | -4.2 | 3.7 |
| Visual2-20_R-Ctx | VentroMedial Visual Area 1 | -4.2 | 3.7 |
| Auditory-07_R-Ctx | Auditory 4 Complex | -4.2 | 3.7 |
| Auditory-12_L-Ctx | ParaBelt Complex | -4.18 | 3.7 |
| Visual2-26_R-Ctx | VentroMedial Visual Area 2 | -4.17 | 3.7 |
| Cingulo-Opercular-39_L-Ctx | Area p32 prime | -4.15 | 3.7 |
| Language-07_R-Ctx | Area STSd posterior | -4.15 | 3.7 |
| Default-71_L-Ctx | ParaHippocampal Area 2 | -4.15 | 3.7 |
| Visual2-02_R-Ctx | Sixth Visual Area | -4.14 | 3.7 |
| Cingulo-Opercular-52_L-Ctx | Area Posterior Insular 1 | -4.12 | 3.7 |
| Language-19_L-Ctx | Auditory 5 Complex | -4.11 | 3.7 |
| Cingulo-Opercular-48_L-Ctx | Frontal OPercular Area 1 | -4.1 | 3.7 |
| Cingulo-Opercular-27_R-Ctx | Para-Insular Area | -4.1 | 3.7 |
| Visual2-47_L-Ctx | VentroMedial Visual Area 1 | -4.05 | 3.7 |

| | | | |
|---|---|---|---|
| Default-64_L-Ctx | ParaHippocampal Area 1 | -4.02 | 3.7 |
| Cingulo-Opercular-44_L-Ctx | Area PFcm | -4.02 | 3.7 |
| Language-06_R-Ctx | Auditory 5 Complex | -4 | 3.7 |
| Visual2-29_L-Ctx | Sixth Visual Area | -3.98 | 3.4 |
| Somatomotor-19_R-Ctx | Frontal OPercular Area 2 | -3.98 | 3.4 |
| Visual2-01_R-Ctx | Medial Superior Temporal Area | -3.97 | 3.4 |
| Default-53_L-Ctx | Area 9 Posterior | -3.97 | 3.4 |
| Auditory-03_R-Ctx | Area TA2 | -3.97 | 3.4 |
| Cingulo-Opercular-04_R-Ctx | Area 5m ventral | -3.96 | 3.4 |
| Cingulo-Opercular-16_R-Ctx | Area 43 | -3.96 | 3.4 |
| Default-25_R-Ctx | ParaHippocampal Area 1 | -3.96 | 3.4 |
| Posterior-Multimodal-02_R-Ctx | Superior Temporal Visual Area | -3.96 | 3.4 |
| Language-02_R-Ctx | Superior Frontal Language Area | -3.96 | 3.4 |
| Default-52_L-Ctx | Area 8B Lateral | -3.93 | 3.4 |
| Auditory-06_R-Ctx | Lateral Belt Complex | -3.93 | 3.4 |
| Default-15_R-Ctx | Area 9 Posterior | -3.93 | 3.4 |
| Language-20_L-Ctx | Area STSd anterior | -3.93 | 3.4 |
| Language-01_R-Ctx | Area 55b | -3.92 | 3.4 |
| Cingulo-Opercular-06_R-Ctx | Supplementary and Cingulate Eye Field | -3.92 | 3.4 |
| Cingulo-Opercular-17_R-Ctx | Area PFcm | -3.91 | 3.4 |
| Default-63_L-Ctx | Hippocampus | -3.9 | 3.4 |
| Posterior-Multimodal-10_L-Cerebellum | | -3.89 | 3.4 |
| Cingulo-Opercular-43_L-Ctx | Area 43 | -3.89 | 3.4 |
| Cingulo-Opercular-10_R-Ctx | Anterior 24 prime | -3.89 | 3.4 |
| Default-37_R-Ctx | Area STSv anterior | -3.88 | 3.4 |
| Visual1-06_L-Ctx | Dorsal Transitional Visual Area | -3.88 | 3.4 |
| Auditory-14_L-Ctx | Lateral Belt Complex | -3.88 | 3.4 |
| Default-14_R-Ctx | Area 8B Lateral | -3.88 | 3.4 |
| Visual2-27_R-Ctx | Ventral Visual Complex | -3.88 | 3.4 |
| Cingulo-Opercular-11_R-Ctx | Area p32 prime | -3.86 | 3.4 |
| Somatomotor-19_R-Hippocampus | | -3.86 | 3.4 |
| Visual2-21_R-Ctx | VentroMedial Visual Area 3 | -3.85 | 3.4 |

| | | | |
|---|---|---|---|
| Default-26_R-Ctx | Area STSd anterior | -3.85 | 3.4 |
| Cingulo-Opercular-19_R-Ctx | Frontal OPercular Area 4 | -3.83 | 3.22 |
| Visual1-03_R-Ctx | Dorsal Transitional Visual Area | -3.83 | 3.22 |
| Visual2-08_L-Cerebellum | | -3.82 | 3.22 |
| Cingulo-Opercular-01_R-Ctx | Frontal Eye Fields | -3.82 | 3.22 |
| Default-18_R-Ctx | Area 9 anterior | -3.82 | 3.22 |
| Somatomotor-18_R-Ctx | RetroInsular Cortex | -3.81 | 3.22 |
| Visual2-28_L-Ctx | Medial Superior Temporal Area | -3.81 | 3.1 |
| Cingulo-Opercular-07_R-Ctx | Area 6m anterior | -3.8 | 3 |
| Somatomotor-28_R-Thalamus | | -3.8 | 3 |
| Somatomotor-13_R-Ctx | Area 6mp | -3.79 | 3 |
| Language-16_L-Ctx | Area IFJa | -3.79 | 3 |
| Language-04_R-Ctx | Area IFJa | -3.79 | 3 |
| Visual2-19_R-Ctx | Area V6A | -3.78 | 3 |
| Language-13_L-Ctx | Superior Temporal Visual Area | -3.78 | 3 |
| Cingulo-Opercular-03_R-Ctx | PeriSylvian Language Area | -3.78 | 3 |
| Visual2-48_L-Ctx | VentroMedial Visual Area 3 | -3.78 | 3 |
| Cingulo-Opercular-45_L-Ctx | Posterior Insular Area 2 | -3.78 | 3 |
| Visual2-46_L-Ctx | Area V6A | -3.77 | 3 |
| Cingulo-Opercular-30_L-Ctx | Frontal Eye Fields | -3.77 | 3 |
| Visual2-42_L-Ctx | Middle Temporal Area | -3.76 | 3 |
| Language-21_L-Ctx | Area STSd posterior | -3.76 | 3 |
| Auditory-10_L-Ctx | RetroInsular Cortex | -3.75 | 3 |
| Cingulo-Opercular-32_L-Ctx | Area 23c | -3.74 | 2.92 |
| Language-08_R-Ctx | Area TemporoParietoOcci pital Junction 1 | -3.74 | 2.92 |
| Visual1-02_R-Ctx | ProStriate Area | -3.73 | 2.92 |
| Somatomotor-38_L-Ctx | Frontal OPercular Area 2 | -3.73 | 2.92 |
| Cingulo-Opercular-31_L-Ctx | Area 5m ventral | -3.71 | 2.85 |
| Visual2-54_L-Ctx | Ventral Visual Complex | -3.71 | 2.85 |
| Visual2-15_R-Cerebellum | | -3.71 | 2.85 |
| Somatomotor-15_R-Ctx | Area OP4/PV | -3.71 | 2.85 |
| Dorsal-Attention-04_R-Ctx | Area 6 anterior | -3.71 | 2.85 |
| Visual1-05_L-Ctx | ProStriate Area | -3.7 | 2.85 |

| | | | |
|---|---|---|---|
| Language-17_L-Ctx | Area IFSp | -3.69 | 2.85 |
| Auditory-02_R-Ctx | Area 52 | -3.68 | 2.8 |
| Somatomotor-08_R-Ctx | Area 7PC | -3.68 | 2.8 |
| Somatomotor-06_R-Ctx | Ventral Area 24d | -3.67 | 2.74 |
| Dorsal-Attention-09_L-Cerebellum | | -3.67 | 2.74 |
| Dorsal-Attention-20_L-Ctx | Area TE2 posterior | -3.66 | 2.7 |
| Somatomotor-14_R-Ctx | Ventral Area 6 | -3.66 | 2.7 |
| Language-10_L-Ctx | Area 55b | -3.66 | 2.7 |
| Language-12_L-Ctx | Superior Frontal Language Area | -3.65 | 2.7 |
| Visual2-22_R-Ctx | Area V4t | -3.65 | 2.7 |
| Dorsal-Attention-16_R-Cerebellum | | -3.63 | 2.7 |
| Visual2-15_R-Ctx | Middle Temporal Area | -3.63 | 2.7 |
| Default-23_L-Hippocampus | | -3.63 | 2.7 |
| Visual2-25_R-Ctx | Area Lateral Occipital 3 | -3.62 | 2.7 |
| Orbito-Affective-03_R-Ctx | posterior OFC Complex | -3.62 | 2.7 |
| Visual2-08_R-Ctx | Seventh Visual Area | -3.61 | 2.7 |
| Frontoparietal-04_R-Ctx | Area 33 prime | -3.61 | 2.7 |
| Cingulo-Opercular-02_R-Ctx | Premotor Eye Field | -3.61 | 2.7 |
| Posterior-Multimodal-05_L-Ctx | PreCuneus Visual Area | -3.61 | 2.7 |
| Ventral-Multimodal-04_L-Ctx | Area TF | -3.61 | 2.7 |
| Somatomotor-28_L-Ctx | Area 7PC | -3.6 | 2.7 |
| Orbito-Affective-01_R-Ctx | Pirform Cortex | -3.6 | 2.7 |
| Cingulo-Opercular-33_L-Ctx | Supplementary and Cingulate Eye Field | -3.59 | 2.7 |
| Posterior-Multimodal-03_R-Ctx | Area TemporoParietoOcci pital Junction 2 | -3.59 | 2.7 |
| Cingulo-Opercular-37_L-Ctx | Area 33 prime | -3.59 | 2.7 |
| Cingulo-Opercular-05_R-Ctx | Area 23c | -3.59 | 2.7 |
| Visual2-38_L-Ctx | Area V3B | -3.59 | 2.7 |
| Cingulo-Opercular-17_R-Cerebellum | | -3.59 | 2.7 |
| Auditory-11_L-Ctx | Area TA2 | -3.58 | 2.62 |
| Default-07_R-Ctx | Area a24 | -3.58 | 2.62 |
| Cingulo-Opercular-56_L-Ctx | Area posterior 24 | -3.58 | 2.62 |
| Visual2-16_R-Ctx | Area Lateral IntraParietal ventral | -3.58 | 2.62 |

| | | | |
|---|---|---|---|
| Posterior-Multimodal-01_R-Ctx | PreCuneus Visual Area | -3.57 | 2.62 |
| Default-76_L-Ctx | Area STSv anterior | -3.57 | 2.62 |
| Posterior-Multimodal-07_L-Ctx | Area TemporoParietoOcci pital Junction 3 | -3.57 | 2.62 |
| Language-11_L-Ctx | PeriSylvian Language Area | -3.56 | 2.55 |
| Visual2-43_L-Ctx | Area Lateral IntraParietal ventral | -3.56 | 2.55 |
| Orbito-Affective-05_L-Ctx | Anterior Agranular Insula Complex | -3.55 | 2.55 |
| Auditory-05_R-Ctx | Medial Belt Complex | -3.55 | 2.55 |
| Somatomotor-26_L-Ctx | Ventral Area 24d | -3.55 | 2.52 |
| Somatomotor-36_L-Ctx | Area OP1/SII | -3.54 | 2.52 |
| Dorsal-Attention-17_L-Ctx | Area PFt | -3.54 | 2.49 |
| Language-05_R-Ctx | Area STGa | -3.54 | 2.49 |
| Cingulo-Opercular-09_R-Ctx | Area Posterior 24 prime | -3.53 | 2.49 |
| Language-15_L-Ctx | Area 45 | -3.53 | 2.49 |
| Visual2-17_R-Ctx | Ventral IntraParietal Complex | -3.53 | 2.42 |
| Posterior-Multimodal-04_R-Ctx | Area TemporoParietoOcci pital Junction 3 | -3.52 | 2.42 |
| Somatomotor-17_R-Ctx | Area OP2-3/VS | -3.51 | 2.38 |
| Somatomotor-33_L-Ctx | Area 6mp | -3.51 | 2.34 |
| Language-03_R-Ctx | Area 45 | -3.51 | 2.34 |
| Visual2-12_R-Ctx | Area Lateral Occipital 1 | -3.51 | 2.3 |
| Cingulo-Opercular-34_L-Ctx | Area 6m anterior | -3.5 | 2.3 |
| Cingulo-Opercular-41_L-Ctx | Area 46 | -3.5 | 2.3 |
| Somatomotor-27_L-Thalamus | | -3.5 | 2.28 |
| Cingulo-Opercular-55_L-Ctx | Area anterior 32 prime | -3.5 | 2.28 |
| Default-44_L-Ctx | Area a24 | -3.5 | 2.28 |
| Visual2-27_R-Hippocampus | | -3.5 | 2.28 |
| Dorsal-Attention-12_L-Ctx | Premotor Eye Field | -3.5 | 2.28 |
| Somatomotor-03_R-Amygdala | | -3.5 | 2.27 |
| Somatomotor-07_R-Ctx | Lateral Area 7A | -3.49 | 2.25 |
| Visual2-37_L-Ctx | Fusiform Face Complex | -3.49 | 2.25 |
| Visual2-06_R-Ctx | Eighth Visual Area | -3.49 | 2.25 |
| Auditory-13_L-Ctx | Medial Belt Complex | -3.49 | 2.25 |
| Visual2-34_L-Ctx | Area V3A | -3.49 | 2.22 |

| | | | |
|---|---|---|---|
| Visual2-09_R-Ctx | IntraParietal Sulcus Area 1 | -3.48 | 2.22 |
| Default-56_L-Ctx | Area 9 anterior | -3.48 | 2.22 |
| Somatomotor-25_L-Ctx | Dorsal Area 24d | -3.48 | 2.21 |
| Default-50_L-Ctx | Area 8Ad | -3.48 | 2.21 |
| Default-55_L-Ctx | Area 47l (47 lateral) | -3.47 | 2.18 |
| Dorsal-Attention-16_L-Ctx | Area 6 anterior | -3.47 | 2.18 |
| Somatomotor-32_L-Ctx | Dorsal area 6 | -3.47 | 2.17 |
| Default-27_R-Ctx | Area STSv posterior | -3.47 | 2.17 |
| Cingulo-Opercular-23_R-Ctx | Area PF opercular | -3.47 | 2.17 |
| Visual2-35_L-Ctx | Seventh Visual Area | -3.46 | 2.17 |
| Somatomotor-16_R-Ctx | Area OP1/SII | -3.46 | 2.17 |
| Somatomotor-12_R-Ctx | Dorsal area 6 | -3.46 | 2.17 |
| Default-47_L-Ctx | Area 10r | -3.46 | 2.17 |
| Ventral-Multimodal-02_R-Ctx | Area TF | -3.45 | 2.15 |
| Somatomotor-34_L-Ctx | Ventral Area 6 | -3.45 | 2.14 |
| Dorsal-Attention-14_L-Ctx | Medial IntraParietal Area | -3.45 | 2.14 |
| Somatomotor-35_L-Ctx | Area OP4/PV | -3.45 | 2.14 |
| Cingulo-Opercular-08_R-Ctx | Medial Area 7A | -3.44 | 2.14 |
| Frontoparietal-35_L-Ctx | Area IFJp | -3.44 | 2.13 |
| Dorsal-Attention-03_R-Ctx | Area Lateral IntraParietal dorsal | -3.43 | 2.11 |
| Language-07_R-Cerebellum | | -3.43 | 2.11 |
| Somatomotor-37_L-Ctx | Area OP2-3/VS | -3.42 | 2.11 |
| Cingulo-Opercular-36_L-Ctx | Area Posterior 24 prime | -3.42 | 2.09 |
| Frontoparietal-10_R-Ctx | Area IFJp | -3.42 | 2.09 |
| Somatomotor-16_L-Hippocampus | | -3.42 | 2.07 |
| Auditory-14_L-Hippocampus | | -3.41 | 2.06 |
| Language-22_L-Ctx | Area TemporoParietoOcci pital Junction 1 | -3.41 | 2.06 |
| Frontoparietal-10_L-Caudate | | -3.41 | 2.06 |
| Visual2-49_L-Ctx | Area V4t | -3.41 | 2.06 |
| Somatomotor-05_R-Ctx | Dorsal Area 24d | -3.41 | 2.06 |
| Dorsal-Attention-05_R-Ctx | Area PFt | -3.41 | 2.03 |
| Visual2-07_R-Ctx | Area V3A | -3.4 | 2.03 |
| Visual2-10_R-Ctx | Fusiform Face Complex | -3.4 | 2 |
| Somatomotor-20_R-Ctx | Insular Granular Complex | -3.39 | 1.99 |
| Language-05_L-Cerebellum | | -3.39 | 1.98 |
| Somatomotor-24_L-Ctx | Area 5L | -3.39 | 1.98 |

| | | | |
|---|---|---|---|
| Dorsal-Attention-22_L-Ctx | Area PGp | -3.39 | 1.97 |
| Language-06_R-Cerebellum | | -3.38 | 1.95 |
| Somatomotor-10_R-Ctx | Area 2 | -3.38 | 1.95 |
| Visual2-36_L-Ctx | IntraParietal Sulcus Area 1 | -3.38 | 1.95 |
| Dorsal-Attention-02_R-Ctx | Medial IntraParietal Area | -3.37 | 1.93 |
| Auditory-01_R-Ctx | Primary Auditory Cortex | -3.37 | 1.93 |
| Posterior-Multimodal-18_R-Cerebellum | | -3.37 | 1.93 |
| Frontoparietal-11_R-Caudate | | -3.37 | 1.93 |
| Somatomotor-03_R-Ctx | Area 5m | -3.37 | 1.93 |
| Default-24_R-Ctx | Hippocampus | -3.37 | 1.91 |
| Dorsal-Attention-18_L-Ctx | Anterior IntraParietal Area | -3.36 | 1.91 |
| Somatomotor-23_L-Ctx | Area 5m | -3.36 | 1.87 |
| Somatomotor-09_R-Ctx | Area 1 | -3.36 | 1.87 |
| Visual2-13_R-Ctx | Area Lateral Occipital 2 | -3.35 | 1.87 |
| Default-12_R-Ctx | Area 8Ad | -3.35 | 1.86 |
| Default-45_L-Ctx | Area dorsal 32 | -3.35 | 1.85 |
| Dorsal-Attention-23_L-Ctx | Area IntraParietal 0 | -3.34 | 1.84 |
| Visual2-51_L-Ctx | Area V3CD | -3.34 | 1.84 |
| Cingulo-Opercular-26_R-Ctx | Area Frontal Opercular 5 | -3.34 | 1.84 |
| Default-51_L-Ctx | Area 9 Middle | -3.33 | 1.84 |
| Default-65_L-Ctx | Area STSv posterior | -3.33 | 1.81 |
| Somatomotor-27_L-Ctx | Lateral Area 7A | -3.33 | 1.8 |
| Visual2-44_L-Ctx | Ventral IntraParietal Complex | -3.33 | 1.8 |
| Cingulo-Opercular-46_L-Ctx | Frontal OPercular Area 4 | -3.32 | 1.79 |
| Visual2-39_L-Ctx | Area Lateral Occipital 1 | -3.31 | 1.77 |
| Dorsal-Attention-13_L-Ctx | Lateral Area 7P | -3.31 | 1.77 |
| Posterior-Multimodal-11_L-Cerebellum | | -3.31 | 1.77 |
| Frontoparietal-36_L-Ctx | Area IFSa | -3.3 | 1.75 |
| Visual2-52_L-Ctx | Area Lateral Occipital 3 | -3.3 | 1.75 |
| Somatomotor-30_L-Ctx | Area 2 | -3.3 | 1.74 |
| Visual2-23_R-Ctx | Area FST | -3.3 | 1.74 |
| Visual2-30_L-Ctx | Second Visual Area | -3.29 | 1.73 |
| Visual2-03_R-Ctx | Second Visual Area | -3.29 | 1.73 |
| Visual2-33_L-Ctx | Eighth Visual Area | -3.28 | 1.73 |
| Dorsal-Attention-01_R-Ctx | Lateral Area 7P | -3.28 | 1.72 |
| Cingulo-Opercular-35_L-Ctx | Medial Area 7A | -3.28 | 1.72 |

| | | | |
|---|---|---|---|
| Visual2-31_L-Ctx | Third Visual Area | -3.27 | 1.69 |
| Somatomotor-01_R-Ctx | Primary Motor Cortex | -3.27 | 1.69 |
| Cingulo-Opercular-28_R-Ctx | Area anterior 32 prime | -3.27 | 1.69 |
| Somatomotor-04_R-Ctx | Area 5L | -3.26 | 1.67 |
| Somatomotor-02_R-Ctx | Primary Sensory Cortex | -3.26 | 1.66 |
| Visual2-10_L-Cerebellum | | -3.25 | 1.65 |
| Dorsal-Attention-10_R-Ctx | Area PGp | -3.25 | 1.64 |
| Somatomotor-21_L-Ctx | Primary Motor Cortex | -3.25 | 1.64 |
| Dorsal-Attention-11_R-Ctx | Area IntraParietal 0 | -3.24 | 1.63 |
| Dorsal-Attention-06_R-Ctx | Anterior IntraParietal Area | -3.24 | 1.61 |
| Visual2-11_R-Ctx | Area V3B | -3.24 | 1.61 |

*Note*. [a]CAB-NP label per Cole-Anticevic Brain Network Parcellation v1.1.6.

[b]Glasser cortical label names were from Glasser, et al. (2016).

[c]*z*-score from Aspin-Welch's test for using exchangeability blocks to adjusted for four scanning sites and variance was estimated for each block, implemented in FSL PALM.

**Supplementary Table S3**

***Age and Nodal Betweenness Centrality Association Results***

| CAB-NP Label[a] | Glasser Label[b] | z[c] | -log10($p_{FWE}$) |
|---|---|---|---|
| Visual1-51_L-Putamen | | 4.45 | 1.91 |
| Somatomotor-25_L-Putamen | | 4.39 | 1.8 |

*Note*. [a]CAB-NP label per Cole-Anticevic Brain Network Parcellation v1.1.6.

[b]Glasser cortical label names were from Glasser, et al. (2016).

[c]*z*-score from Aspin-Welch's test for using exchangeability blocks to adjusted for four scanning sites and variance was estimated for each block, implemented in FSL PALM.

**Supplementary Table S4**

***Age and Nodal Eigenvector Centrality Association Results***

| CAB-NP Label[a] | Glasser Label[b] | z[c] | -log10($p_{FWE}$) |
|---|---|---|---|
| | Positive Association | | |
| Frontoparietal-27_R-Cerebellum | | 4.57 | 3.7 |
| Frontoparietal-24_R-Ctx | Area PFm Complex | 4.55 | 3.7 |
| Frontoparietal-48_L-Ctx | Area PFm Complex | 4.39 | 3.7 |
| Frontoparietal-15_R-Ctx | Area 11l | 4.24 | 3.7 |
| Frontoparietal-40_L-Ctx | Area 11l | 4.2 | 3.7 |
| Frontoparietal-13_L-Cerebellum | | 4.13 | 3.7 |
| Somatomotor-22_L-Pallidum | | 4.11 | 3.7 |
| Frontoparietal-23_R-Ctx | Area IntraParietal 1 | 4.1 | 3.7 |
| Cingulo-Opercular-25_L-Diencephalon | | 4.09 | 3.7 |
| Frontoparietal-37_R-Cerebellum | | 4.07 | 3.7 |
| Frontoparietal-07_R-Ctx | Area 8C | 4.04 | 3.7 |
| Frontoparietal-09_R-Ctx | Area anterior 47r | 3.98 | 3.7 |
| Auditory-06_L-Cerebellum | | 3.95 | 3.7 |
| Frontoparietal-39_L-Diencephalon | | 3.95 | 3.7 |
| Frontoparietal-46_L-Ctx | Area IntraParietal 2 | 3.95 | 3.7 |
| Default-03_R-Ctx | Area 23d | 3.95 | 3.7 |
| Frontoparietal-30_L-Ctx | Parieto-Occipital Sulcus Area 2 | 3.89 | 3.7 |
| Visual1-47_L-Pallidum | | 3.87 | 3.7 |
| Default-40_L-Ctx | Area 23d | 3.84 | 3.4 |
| Visual1-11_L-Brainstem | | 3.84 | 3.4 |
| Visual1-48_R-Pallidum | | 3.83 | 3.22 |
| Frontoparietal-33_L-Ctx | Area 8C | 3.83 | 3.22 |
| Frontoparietal-04_R-Brainstem | | 3.83 | 3.22 |
| Default-21_R-Diencephalon | | 3.82 | 3.22 |
| Visual1-28_L-Cerebellum | | 3.82 | 3.1 |
| Auditory-31_R-Thalamus | | 3.82 | 3 |
| Auditory-20_L-Pallidum | | 3.8 | 3 |
| Frontoparietal-22_R-Ctx | Area IntraParietal 2 | 3.78 | 2.85 |
| Ventral-Multimodal-04_R-Hippocampus | | 3.76 | 2.8 |
| Posterior-Multimodal-05_L-Brainstem | | 3.75 | 2.74 |
| Default-17_R-Cerebellum | | 3.74 | 2.66 |
| Default-32_R-Ctx | Area PGs | 3.73 | 2.62 |

| | | | |
|---|---|---|---|
| Dorsal-Attention-01_L-Amygdala | | 3.7 | 2.62 |
| Frontoparietal-12_R-Ctx | Area posterior 9-46v | 3.66 | 2.44 |
| Somatomotor-15_L-Diencephalon | | 3.61 | 2.34 |
| Frontoparietal-47_L-Ctx | Area IntraParietal 1 | 3.59 | 2.3 |
| Auditory-08_L-Diencephalon | | 3.59 | 2.3 |
| Visual1-49_R-Pallidum | | 3.58 | 2.27 |
| Frontoparietal-50_L-Ctx | Area posterior 47r | 3.58 | 2.24 |
| Frontoparietal-19_L-Cerebellum | | 3.58 | 2.24 |
| Auditory-11_R-Diencephalon | | 3.55 | 2.15 |
| Orbito-Affective-12_R-Pallidum | | 3.55 | 2.15 |
| Frontoparietal-36_R-Cerebellum | | 3.53 | 2.08 |
| Visual1-02_L-Accumbens | | 3.52 | 2.03 |
| Frontoparietal-47_L-Thalamus | | 3.52 | 2.02 |
| Frontoparietal-45_L-Ctx | Area TE1 posterior | 3.5 | 1.93 |
| Orbito-Affective-03_L-Caudate | | 3.49 | 1.89 |
| Frontoparietal-34_L-Ctx | Area anterior 47r | 3.48 | 1.82 |
| Default-16_R-Cerebellum | | 3.47 | 1.8 |
| Auditory-07_R-Cerebellum | | 3.45 | 1.75 |
| Cingulo-Opercular-06_L-Brainstem | | 3.44 | 1.7 |
| Frontoparietal-42_L-Pallidum | | 3.44 | 1.7 |
| Cingulo-Opercular-26_L-Diencephalon | | 3.42 | 1.66 |
| | Negative Association | | |
| Visual2-20_R-Ctx | VentroMedial Visual Area 1 | -4.8 | 3.7 |
| Visual2-02_R-Ctx | Sixth Visual Area | -4.64 | 3.7 |
| Auditory-07_R-Ctx | Auditory 4 Complex | -4.55 | 3.7 |
| Auditory-15_L-Ctx | Auditory 4 Complex | -4.38 | 3.7 |
| Auditory-04_R-Ctx | ParaBelt Complex | -4.31 | 3.7 |
| Visual2-26_R-Ctx | VentroMedial Visual Area 2 | -4.3 | 3.7 |
| Auditory-12_L-Ctx | ParaBelt Complex | -4.3 | 3.7 |
| Visual2-29_L-Ctx | Sixth Visual Area | -4.16 | 3.7 |
| Visual2-47_L-Ctx | VentroMedial Visual Area 1 | -4.14 | 3.7 |
| Cingulo-Opercular-18_R-Ctx | Posterior Insular Area 2 | -4.03 | 3.7 |
| Visual2-53_L-Ctx | VentroMedial Visual Area 2 | -3.97 | 3.7 |
| Auditory-03_R-Ctx | Area TA2 | -3.83 | 3.22 |

| | | | |
|---|---|---|---|
| Auditory-06_R-Ctx | Lateral Belt Complex | -3.82 | 3 |
| Language-06_R-Ctx | Auditory 5 Complex | -3.79 | 2.92 |
| Cingulo-Opercular-04_R-Ctx | Area 5m ventral | -3.76 | 2.74 |
| Auditory-14_L-Ctx | Lateral Belt Complex | -3.73 | 2.62 |
| Cingulo-Opercular-44_L-Ctx | Area PFcm | -3.73 | 2.62 |
| Visual2-01_R-Ctx | Medial Superior Temporal Area | -3.72 | 2.62 |
| Somatomotor-18_R-Ctx | RetroInsular Cortex | -3.64 | 2.42 |
| Auditory-10_L-Ctx | RetroInsular Cortex | -3.55 | 2.15 |
| Visual1-06_L-Ctx | Dorsal Transitional Visual Area | -3.54 | 2.12 |
| Visual1-03_R-Ctx | Dorsal Transitional Visual Area | -3.52 | 2.03 |
| Visual1-02_R-Ctx | ProStriate Area | -3.5 | 1.94 |
| Cingulo-Opercular-21_R-Ctx | Frontal OPercular Area 1 | -3.5 | 1.92 |
| Language-19_L-Ctx | Auditory 5 Complex | -3.49 | 1.86 |
| Dorsal-Attention-07_R-Ctx | ParaHippocampal Area 3 | -3.48 | 1.84 |
| Default-33_R-Ctx | ParaHippocampal Area 2 | -3.48 | 1.81 |
| Visual2-27_R-Ctx | Ventral Visual Complex | -3.46 | 1.79 |
| Visual1-05_L-Ctx | ProStriate Area | -3.44 | 1.7 |

*Note*. [a]CAB-NP label per Cole-Anticevic Brain Network Parcellation v1.1.6.

[b]Glasser cortical label names were from Glasser, et al. (2016).

[c]*z*-score from Aspin-Welch's test for using exchangeability blocks to adjusted for four scanning sites and variance was estimated for each block, implemented in FSL PALM.

**Supplementary Table S5**

***Age and Nodal Clustering Coefficient Association Results***

| CAB-NP Label[a] | Glasser Label[b] | z[c] | -log10($p_{FWE}$) |
|---|---|---|---|
| Visual2-02_R-Ctx | Sixth Visual Area | -4.61 | 3.7 |
| Auditory-15_L-Ctx | Auditory 4 Complex | -4.45 | 3.7 |
| Visual2-29_L-Ctx | Sixth Visual Area | -4.42 | 3.7 |
| Cingulo-Opercular-47_L-Ctx | Middle Insular Area | -4.42 | 3.7 |
| Visual2-01_R-Ctx | Medial Superior Temporal Area | -4.4 | 3.7 |
| Auditory-07_R-Ctx | Auditory 4 Complex | -4.39 | 3.7 |
| Cingulo-Opercular-20_R-Ctx | Middle Insular Area | -4.37 | 3.7 |
| Visual2-20_R-Ctx | VentroMedial Visual Area 1 | -4.34 | 3.7 |
| Default-33_R-Ctx | ParaHippocampal Area 2 | -4.31 | 3.7 |
| Cingulo-Opercular-18_R-Ctx | Posterior Insular Area 2 | -4.31 | 3.7 |
| Auditory-04_R-Ctx | ParaBelt Complex | -4.29 | 3.7 |
| Cingulo-Opercular-38_L-Ctx | Anterior 24 prime | -4.27 | 3.7 |
| Cingulo-Opercular-21_R-Ctx | Frontal OPercular Area 1 | -4.27 | 3.7 |
| Auditory-12_L-Ctx | ParaBelt Complex | -4.27 | 3.7 |
| Visual2-53_L-Ctx | VentroMedial Visual Area 2 | -4.24 | 3.7 |
| Cingulo-Opercular-25_R-Ctx | Area Posterior Insular 1 | -4.23 | 3.7 |
| Cingulo-Opercular-22_R-Ctx | Frontal OPercular Area 3 | -4.23 | 3.7 |
| Visual2-28_L-Ctx | Medial Superior Temporal Area | -4.21 | 3.7 |
| Language-19_L-Ctx | Auditory 5 Complex | -4.21 | 3.7 |
| Visual2-46_L-Ctx | Area V6A | -4.2 | 3.7 |
| Visual2-47_L-Ctx | VentroMedial Visual Area 1 | -4.19 | 3.7 |
| Language-06_R-Ctx | Auditory 5 Complex | -4.17 | 3.7 |
| Cingulo-Opercular-04_R-Ctx | Area 5m ventral | -4.14 | 3.7 |
| Visual2-42_L-Ctx | Middle Temporal Area | -4.14 | 3.7 |
| Default-25_R-Ctx | ParaHippocampal Area 1 | -4.13 | 3.7 |
| Dorsal-Attention-07_R-Ctx | ParaHippocampal Area 3 | -4.13 | 3.7 |
| Dorsal-Attention-19_L-Ctx | ParaHippocampal Area 3 | -4.1 | 3.7 |
| Auditory-03_R-Ctx | Area TA2 | -4.1 | 3.7 |

| | | | |
|---|---|---|---|
| Visual2-22_R-Ctx | Area V4t | -4.1 | 3.7 |
| Visual2-26_R-Ctx | VentroMedial Visual Area 2 | -4.09 | 3.7 |
| Visual2-08_R-Ctx | Seventh Visual Area | -4.08 | 3.4 |
| Cingulo-Opercular-39_L-Ctx | Area p32 prime | -4.06 | 3.4 |
| Visual2-15_R-Ctx | Middle Temporal Area | -4.06 | 3.4 |
| Cingulo-Opercular-44_L-Ctx | Area PFcm | -4.04 | 3.4 |
| Visual2-19_R-Ctx | Area V6A | -4.04 | 3.4 |
| Visual1-06_L-Ctx | Dorsal Transitional Visual Area | -4.02 | 3.22 |
| Cingulo-Opercular-52_L-Ctx | Area Posterior Insular 1 | -4.01 | 3.22 |
| Default-64_L-Ctx | ParaHippocampal Area 1 | -4.01 | 3.22 |
| Somatomotor-13_R-Ctx | Area 6mp | -4.01 | 3.22 |
| Visual1-03_R-Ctx | Dorsal Transitional Visual Area | -4 | 3.22 |
| Somatomotor-08_R-Ctx | Area 7PC | -3.99 | 3.22 |
| Visual2-34_L-Ctx | Area V3A | -3.96 | 3.22 |
| Cingulo-Opercular-27_R-Ctx | Para-Insular Area | -3.96 | 3.22 |
| Language-20_L-Ctx | Area STSd anterior | -3.95 | 3.22 |
| Visual2-27_R-Ctx | Ventral Visual Complex | -3.94 | 3.22 |
| Visual2-16_R-Ctx | Area Lateral IntraParietal ventral | -3.92 | 3.22 |
| Cingulo-Opercular-01_R-Ctx | Frontal Eye Fields | -3.92 | 3.22 |
| Auditory-14_L-Ctx | Lateral Belt Complex | -3.92 | 3.22 |
| Auditory-06_R-Ctx | Lateral Belt Complex | -3.92 | 3.1 |
| Visual2-54_L-Ctx | Ventral Visual Complex | -3.9 | 3.1 |
| Cingulo-Opercular-48_L-Ctx | Frontal OPercular Area 1 | -3.9 | 3.1 |
| Cingulo-Opercular-16_R-Ctx | Area 43 | -3.9 | 3 |
| Cingulo-Opercular-17_R-Ctx | Area PFcm | -3.89 | 3 |
| Orbito-Affective-01_R-Ctx | Pirform Cortex | -3.89 | 3 |
| Visual1-02_R-Ctx | ProStriate Area | -3.89 | 3 |
| Language-13_L-Ctx | Superior Temporal Visual Area | -3.89 | 3 |
| Visual2-38_L-Ctx | Area V3B | -3.89 | 3 |
| Default-71_L-Ctx | ParaHippocampal Area 2 | -3.88 | 3 |
| Cingulo-Opercular-49_L-Ctx | Frontal OPercular Area 3 | -3.88 | 3 |
| Language-07_R-Ctx | Area STSd posterior | -3.87 | 3 |

| | | | |
|---|---|---|---|
| Somatomotor-19_R-Ctx | Frontal OPercular Area 2 | -3.85 | 3 |
| Visual2-15_R-Cerebellum | | -3.84 | 3 |
| Cingulo-Opercular-06_R-Ctx | Supplementary and Cingulate Eye Field | -3.84 | 3 |
| Posterior-Multimodal-02_R-Ctx | Superior Temporal Visual Area | -3.83 | 2.92 |
| Visual2-25_R-Ctx | Area Lateral Occipital 3 | -3.83 | 2.92 |
| Somatomotor-28_L-Ctx | Area 7PC | -3.83 | 2.92 |
| Cingulo-Opercular-43_L-Ctx | Area 43 | -3.83 | 2.92 |
| Visual2-35_L-Ctx | Seventh Visual Area | -3.83 | 2.92 |
| Dorsal-Attention-04_R-Ctx | Area 6 anterior | -3.83 | 2.92 |
| Visual2-43_L-Ctx | Area Lateral IntraParietal ventral | -3.83 | 2.92 |
| Somatomotor-28_R-Thalamus | | -3.81 | 2.92 |
| Default-63_L-Ctx | Hippocampus | -3.81 | 2.92 |
| Somatomotor-12_R-Ctx | Dorsal area 6 | -3.81 | 2.92 |
| Visual1-05_L-Ctx | ProStriate Area | -3.81 | 2.92 |
| Visual2-17_R-Ctx | Ventral IntraParietal Complex | -3.81 | 2.92 |
| Visual2-48_L-Ctx | VentroMedial Visual Area 3 | -3.8 | 2.92 |
| Cingulo-Opercular-30_L-Ctx | Frontal Eye Fields | -3.79 | 2.92 |
| Cingulo-Opercular-31_L-Ctx | Area 5m ventral | -3.79 | 2.92 |
| Posterior-Multimodal-03_R-Ctx | Area TemporoParietoOcci pital Junction 2 | -3.79 | 2.92 |
| Visual2-07_R-Ctx | Area V3A | -3.79 | 2.92 |
| Default-18_R-Ctx | Area 9 anterior | -3.77 | 2.92 |
| Auditory-11_L-Ctx | Area TA2 | -3.77 | 2.92 |
| Somatomotor-09_R-Ctx | Area 1 | -3.77 | 2.92 |
| Somatomotor-15_R-Ctx | Area OP4/PV | -3.77 | 2.92 |
| Somatomotor-33_L-Ctx | Area 6mp | -3.76 | 2.92 |
| Visual2-49_L-Ctx | Area V4t | -3.76 | 2.85 |
| Cingulo-Opercular-45_L-Ctx | Posterior Insular Area 2 | -3.75 | 2.8 |
| Somatomotor-05_R-Ctx | Dorsal Area 24d | -3.75 | 2.8 |
| Somatomotor-19_R-Hippocampus | | -3.74 | 2.74 |
| Posterior-Multimodal-07_L-Ctx | Area TemporoParietoOcci pital Junction 3 | -3.74 | 2.74 |
| Visual2-21_R-Ctx | VentroMedial Visual Area 3 | -3.74 | 2.74 |

| | | | |
|---|---|---|---|
| Cingulo-Opercular-10_R-Ctx | Anterior 24 prime | -3.73 | 2.74 |
| Visual2-09_R-Ctx | IntraParietal Sulcus Area 1 | -3.73 | 2.7 |
| Visual2-36_L-Ctx | IntraParietal Sulcus Area 1 | -3.73 | 2.7 |
| Somatomotor-23_L-Ctx | Area 5m | -3.72 | 2.7 |
| Somatomotor-36_L-Ctx | Area OP1/SII | -3.72 | 2.7 |
| Visual2-08_L-Cerebellum | | -3.72 | 2.7 |
| Somatomotor-18_R-Ctx | RetroInsular Cortex | -3.72 | 2.7 |
| Somatomotor-06_R-Ctx | Ventral Area 24d | -3.72 | 2.7 |
| Somatomotor-32_L-Ctx | Dorsal area 6 | -3.71 | 2.7 |
| Posterior-Multimodal-04_R-Ctx | Area TemporoParietoOcci pital Junction 3 | -3.71 | 2.7 |
| Dorsal-Attention-09_L-Cerebellum | | -3.69 | 2.7 |
| Visual2-52_L-Ctx | Area Lateral Occipital 3 | -3.69 | 2.7 |
| Somatomotor-26_L-Ctx | Ventral Area 24d | -3.69 | 2.7 |
| Somatomotor-24_L-Ctx | Area 5L | -3.68 | 2.66 |
| Visual2-30_L-Ctx | Second Visual Area | -3.68 | 2.66 |
| Somatomotor-07_R-Ctx | Lateral Area 7A | -3.67 | 2.62 |
| Cingulo-Opercular-11_R-Ctx | Area p32 prime | -3.67 | 2.62 |
| Somatomotor-03_R-Ctx | Area 5m | -3.65 | 2.62 |
| Visual2-37_L-Ctx | Fusiform Face Complex | -3.65 | 2.62 |
| Auditory-10_L-Ctx | RetroInsular Cortex | -3.64 | 2.59 |
| Cingulo-Opercular-03_R-Ctx | PeriSylvian Language Area | -3.63 | 2.59 |
| Visual2-44_L-Ctx | Ventral IntraParietal Complex | -3.63 | 2.59 |
| Dorsal-Attention-14_L-Ctx | Medial IntraParietal Area | -3.63 | 2.59 |
| Cingulo-Opercular-32_L-Ctx | Area 23c | -3.63 | 2.55 |
| Somatomotor-02_R-Ctx | Primary Sensory Cortex | -3.62 | 2.55 |
| Default-37_R-Ctx | Area STSv anterior | -3.62 | 2.52 |
| Cingulo-Opercular-17_R-Cerebellum | | -3.62 | 2.52 |
| Language-16_L-Ctx | Area IFJa | -3.61 | 2.52 |
| Default-26_R-Ctx | Area STSd anterior | -3.61 | 2.47 |
| Language-01_R-Ctx | Area 55b | -3.61 | 2.44 |
| Visual2-06_R-Ctx | Eighth Visual Area | -3.6 | 2.44 |
| Cingulo-Opercular-23_R-Ctx | Area PF opercular | -3.6 | 2.44 |
| Somatomotor-16_R-Ctx | Area OP1/SII | -3.59 | 2.44 |
| Somatomotor-10_R-Ctx | Area 2 | -3.59 | 2.44 |
| Somatomotor-29_L-Ctx | Area 1 | -3.59 | 2.44 |

| | | | |
|---|---|---|---|
| Dorsal-Attention-20_L-Ctx | Area TE2 posterior | -3.59 | 2.42 |
| Somatomotor-25_L-Ctx | Dorsal Area 24d | -3.59 | 2.42 |
| Somatomotor-04_R-Ctx | Area 5L | -3.58 | 2.42 |
| Language-08_R-Ctx | Area TemporoParietoOcci pital Junction 1 | -3.57 | 2.38 |
| Visual2-03_R-Ctx | Second Visual Area | -3.57 | 2.36 |
| Somatomotor-27_L-Ctx | Lateral Area 7A | -3.57 | 2.36 |
| Dorsal-Attention-16_L-Ctx | Area 6 anterior | -3.57 | 2.36 |
| Visual2-10_R-Ctx | Fusiform Face Complex | -3.57 | 2.36 |
| Visual2-12_R-Ctx | Area Lateral Occipital 1 | -3.57 | 2.34 |
| Language-05_R-Ctx | Area STGa | -3.56 | 2.34 |
| Posterior-Multimodal-10_L-Cerebellum | | -3.56 | 2.34 |
| Default-76_L-Ctx | Area STSv anterior | -3.55 | 2.34 |
| Visual2-31_L-Ctx | Third Visual Area | -3.55 | 2.32 |
| Dorsal-Attention-23_L-Ctx | Area IntraParietal 0 | -3.55 | 2.32 |
| Somatomotor-30_L-Ctx | Area 2 | -3.55 | 2.32 |
| Somatomotor-14_R-Ctx | Ventral Area 6 | -3.54 | 2.32 |
| Visual2-23_R-Ctx | Area FST | -3.54 | 2.3 |
| Language-02_R-Ctx | Superior Frontal Language Area | -3.53 | 2.28 |
| Somatomotor-17_R-Ctx | Area OP2-3/VS | -3.53 | 2.27 |
| Somatomotor-22_L-Ctx | Primary Sensory Cortex | -3.52 | 2.27 |
| Auditory-02_R-Ctx | Area 52 | -3.52 | 2.27 |
| Language-21_L-Ctx | Area STSd posterior | -3.52 | 2.27 |
| Somatomotor-01_R-Ctx | Primary Motor Cortex | -3.52 | 2.27 |
| Dorsal-Attention-12_L-Ctx | Premotor Eye Field | -3.52 | 2.27 |
| Cingulo-Opercular-09_R-Ctx | Area Posterior 24 prime | -3.51 | 2.25 |
| Somatomotor-21_L-Ctx | Primary Motor Cortex | -3.5 | 2.21 |
| Somatomotor-27_L-Thalamus | | -3.5 | 2.21 |
| Visual2-39_L-Ctx | Area Lateral Occipital 1 | -3.5 | 2.18 |
| Dorsal-Attention-02_R-Ctx | Medial IntraParietal Area | -3.5 | 2.15 |
| Default-15_R-Ctx | Area 9 Posterior | -3.49 | 2.15 |
| Visual2-13_R-Ctx | Area Lateral Occipital 2 | -3.49 | 2.15 |
| Default-47_L-Ctx | Area 10r | -3.49 | 2.15 |
| Visual2-51_L-Ctx | Area V3CD | -3.49 | 2.13 |
| Language-17_L-Ctx | Area IFSp | -3.49 | 2.13 |

| | | | |
|---|---|---|---|
| Ventral-Multimodal-04_L-Ctx | Area TF | -3.48 | 2.1 |
| Default-24_R-Ctx | Hippocampus | -3.48 | 2.1 |
| Cingulo-Opercular-07_R-Ctx | Area 6m anterior | -3.48 | 2.09 |
| Cingulo-Opercular-37_L-Ctx | Area 33 prime | -3.48 | 2.08 |
| Visual2-50_L-Ctx | Area FST | -3.47 | 2.05 |
| Somatomotor-38_L-Ctx | Frontal OPercular Area 2 | -3.46 | 2.02 |
| Language-10_L-Ctx | Area 55b | -3.46 | 2.02 |
| Visual2-11_R-Ctx | Area V3B | -3.45 | 1.97 |
| Default-23_L-Hippocampus | | -3.45 | 1.95 |
| Posterior-Multimodal-01_R-Ctx | PreCuneus Visual Area | -3.44 | 1.93 |
| Dorsal-Attention-03_R-Ctx | Area Lateral IntraParietal dorsal | -3.43 | 1.92 |
| Visual2-33_L-Ctx | Eighth Visual Area | -3.42 | 1.9 |
| Orbito-Affective-04_L-Ctx | Pirform Cortex | -3.41 | 1.89 |
| Cingulo-Opercular-33_L-Ctx | Supplementary and Cingulate Eye Field | -3.41 | 1.89 |
| Cingulo-Opercular-36_L-Ctx | Area Posterior 24 prime | -3.4 | 1.88 |
| Posterior-Multimodal-05_L-Ctx | PreCuneus Visual Area | -3.4 | 1.87 |
| Dorsal-Attention-11_R-Ctx | Area IntraParietal 0 | -3.4 | 1.86 |
| Dorsal-Attention-17_L-Ctx | Area PFt | -3.4 | 1.84 |
| Cingulo-Opercular-08_R-Ctx | Medial Area 7A | -3.39 | 1.84 |
| Cingulo-Opercular-02_R-Ctx | Premotor Eye Field | -3.38 | 1.8 |
| Somatomotor-35_L-Ctx | Area OP4/PV | -3.38 | 1.8 |
| Language-04_R-Ctx | Area IFJa | -3.38 | 1.8 |
| Cingulo-Opercular-05_R-Ctx | Area 23c | -3.38 | 1.8 |
| Somatomotor-31_L-Ctx | Area 3a | -3.38 | 1.8 |
| Language-22_L-Ctx | Area TemporoParietoOcci pital Junction 1 | -3.38 | 1.79 |
| Somatomotor-37_L-Ctx | Area OP2-3/VS | -3.37 | 1.75 |
| Dorsal-Attention-16_R-Cerebellum | | -3.35 | 1.72 |
| Visual2-05_R-Ctx | Fourth Visual Area | -3.35 | 1.72 |
| Frontoparietal-04_R-Ctx | Area 33 prime | -3.35 | 1.69 |
| Visual2-27_R-Hippocampus | | -3.34 | 1.69 |
| Dorsal-Attention-18_L-Ctx | Anterior IntraParietal Area | -3.34 | 1.69 |
| Dorsal-Attention-22_L-Ctx | Area PGp | -3.34 | 1.69 |
| Visual2-10_L-Cerebellum | | -3.34 | 1.69 |

| | | | |
|---|---|---|---|
| Somatomotor-03_R-Amygdala | | -3.34 | 1.69 |
| Default-53_L-Ctx | Area 9 Posterior | -3.34 | 1.67 |
| Somatomotor-20_R-Ctx | Insular Granular Complex | -3.33 | 1.67 |
| Visual2-04_R-Ctx | Third Visual Area | -3.32 | 1.67 |
| Cingulo-Opercular-56_L-Ctx | Area posterior 24 | -3.31 | 1.63 |
| Ventral-Multimodal-02_R-Ctx | Area TF | -3.31 | 1.62 |
| Default-62_L-Ctx | PreSubiculum | -3.31 | 1.62 |
| Default-14_R-Ctx | Area 8B Lateral | -3.31 | 1.62 |
| Dorsal-Attention-13_L-Ctx | Lateral Area 7P | -3.3 | 1.61 |
| Dorsal-Attention-05_R-Ctx | Area PFt | -3.3 | 1.61 |
| Visual2-32_L-Ctx | Fourth Visual Area | -3.29 | 1.6 |

*Note*. [a]CAB-NP label per Cole-Anticevic Brain Network Parcellation v1.1.6.

[b]Glasser cortical label names were from Glasser, et al. (2016).

[c]*z*-score from Aspin-Welch's test for using exchangeability blocks to adjusted for four scanning sites and variance was estimated for each block, implemented in FSL PALM.

**Supplementary Table S6**

***Age and Nodal Participation Coefficient Association Results***

| CAB-NP Label[a] | Glasser Label[b] | z[c] | -log10($p_{FWE}$) |
|---|---|---|---|
| | Positive Association | | |
| Cingulo-Opercular-47_L-Ctx | Middle Insular Area | 3.93 | 3.22 |
| Cingulo-Opercular-20_R-Ctx | Middle Insular Area | 3.93 | 3.22 |
| Cingulo-Opercular-49_L-Ctx | Frontal OPercular Area 3 | 3.88 | 3.1 |
| Cingulo-Opercular-11_R-Ctx | Area p32 prime | 3.83 | 2.85 |
| Somatomotor-18_L-Hippocampus | | 3.77 | 2.85 |
| Cingulo-Opercular-22_R-Ctx | Frontal OPercular Area 3 | 3.69 | 2.47 |
| Somatomotor-28_R-Thalamus | | 3.65 | 2.34 |
| Cingulo-Opercular-46_L-Ctx | Frontal OPercular Area 4 | 3.65 | 2.32 |
| Cingulo-Opercular-19_R-Ctx | Frontal OPercular Area 4 | 3.65 | 2.32 |
| Somatomotor-16_L-Hippocampus | | 3.64 | 2.3 |
| Somatomotor-27_L-Thalamus | | 3.6 | 2.19 |
| Cingulo-Opercular-07_R-Ctx | Area 6m anterior | 3.52 | 1.84 |
| Visual2-20_R-Ctx | VentroMedial Visual Area 1 | 3.47 | 1.62 |
| Auditory-09_L-Ctx | Area 52 | 3.46 | 1.61 |
| | Negative Association | | |
| Visual1-56_L-Thalamus | | -3.78 | 2.85 |
| Visual1-02_R-Ctx | ProStriate Area | -3.62 | 2.24 |

*Note*. [a]CAB-NP label per Cole-Anticevic Brain Network Parcellation v1.1.6.

[b]Glasser cortical label names were from Glasser, et al. (2016).

[c]*z*-score from Aspin-Welch's test for using exchangeability blocks to adjusted for four scanning sites and variance was estimated for each block, implemented in FSL PALM.

**Supplementary Table S7**

***Summary of Age and Graph-Based Brain Functional Connectivity Results: Positive-Only Connectivity Weights***

| Graph Measure | Positive Association | | | Negative Association | | |
|---|---|---|---|---|---|---|
| | Cortical | Subcortical | CAB-NP Networks | Cortical | Subcortical | CAB-NP Networks |
| Normalized Strength | -- | -- | -- | 125 | 5 | AUD, CON, DAN, DMN, LAN, ORA, PMM, SMN, VIS1, VIS2 |
| Closeness Centrality | -- | -- | -- | 231 | 26 | Across all 12 networks |
| Betweenness Centrality | 1 | 2 | DMN, SMN, VIS2 | -- | -- | -- |
| Eigenvector Centrality | 23 | 39 | AUD, CON, DAN, DMN, FPN, ORA, SMN, VIS1 | 29 | 1 | AUD, CON, DMN, LAN, SMN, VIS1, VIS2 |
| Clustering Coefficient | -- | -- | -- | 197 | 17 | AUD, CON, DAN, DMN, LAN, ORA, PMM, SMN, VIS1, VIS2, VMN |
| Participation Coefficient | 1 | 4 | CON, SMN | -- | -- | -- |

*Note*. 12 CAB-NP Networks: AUD: Auditory Network; CON: Cingulo-Opercular Network; DAN: Dorsal Attention Network; DMN: Default Mode Network; FPN: Frontoparietal Network; LAN: Language Network; ORA: Orbito-Affective Network; PMM: Posterior Multimodal Network; SMN: Somatomotor Network; VIS1: Primary Visual Network; VIS2: Secondary Visual Network; VMM: Ventral Multimodal Network [3].

**Supplementary Table S8**

***Summary of Age and Graph-Based Brain Functional Connectivity Results: Negative-Only Connectivity Weights***

| Graph Measure | Positive Association | | | Negative Association | | |
|---|---|---|---|---|---|---|
| | Cortical | Subcortical | CAB-NP Networks | Cortical | Subcortical | CAB-NP Networks |
| Normalized Strength | -- | -- | -- | -- | 2 | DMN, FPN |
| Closeness Centrality | -- | -- | -- | 10 | -- | CON, DAN, DMN, LAN, ORA, VIS2 |
| Betweenness Centrality | -- | -- | -- | -- | -- | -- |
| Eigenvector Centrality | -- | 1 | VIS1 | -- | 1 | FPN |
| Clustering Coefficient | -- | -- | -- | -- | -- | -- |
| Participation Coefficient | -- | -- | -- | 1 | 1 | SMN, VIS2 |

*Note*. 12 CAB-NP Networks: AUD: Auditory Network; CON: Cingulo-Opercular Network; DAN: Dorsal Attention Network; DMN: Default Mode Network; FPN: Frontoparietal Network; LAN: Language Network; ORA: Orbito-Affective Network; PMM: Posterior Multimodal Network; SMN: Somatomotor Network; VIS1: Primary Visual Network; VIS2: Secondary Visual Network; VMM: Ventral Multimodal Network [3].

**Supplementary Figure S1**

*Age and Graph-Based Brain Functional Connectivity Association Maps: Positive-Only Connectivity Weights. Color bar shows the z-score from Aspin-Welch's test for using exchangeability blocks to adjusted for four scanning sites and variance was estimated for each block, implemented in FSL PALM.*

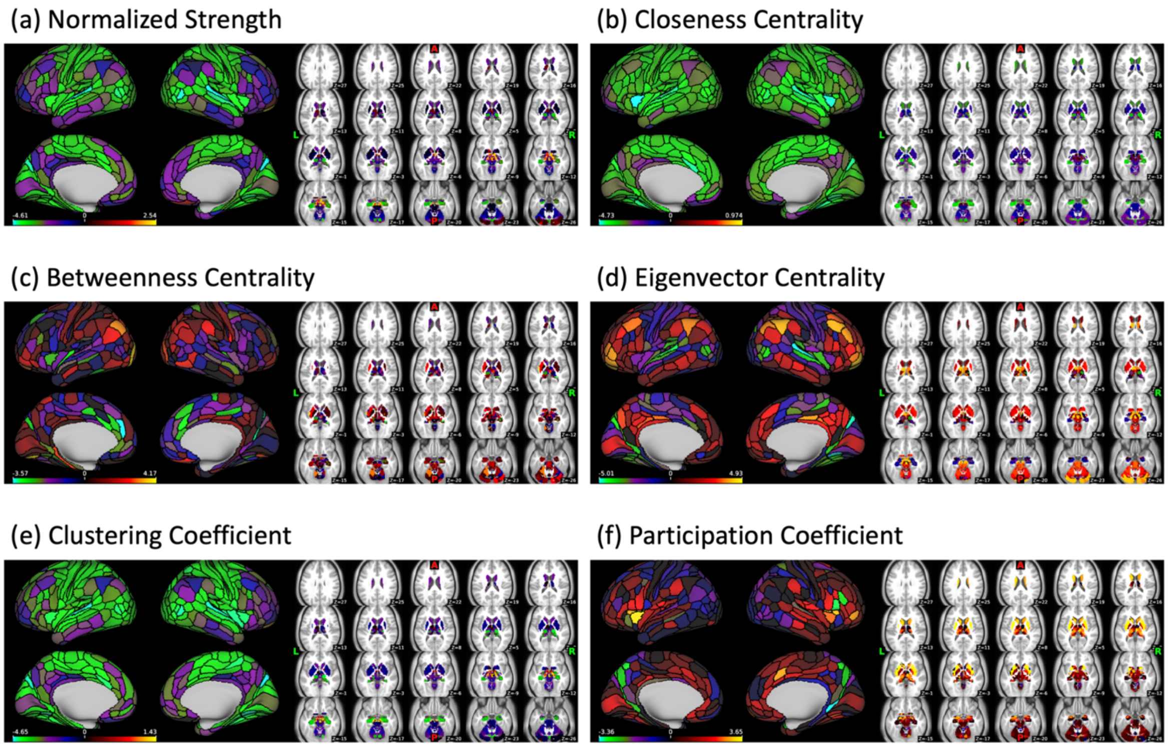

**Supplementary Figure S2**

*Association of Age and Global Graph-Based Brain Functional Connectivity: Positive-Only Connectivity Weights. Modularity association with (a) age and (b) loneliness. Average shortest path length with (c) age and (d) loneliness.*

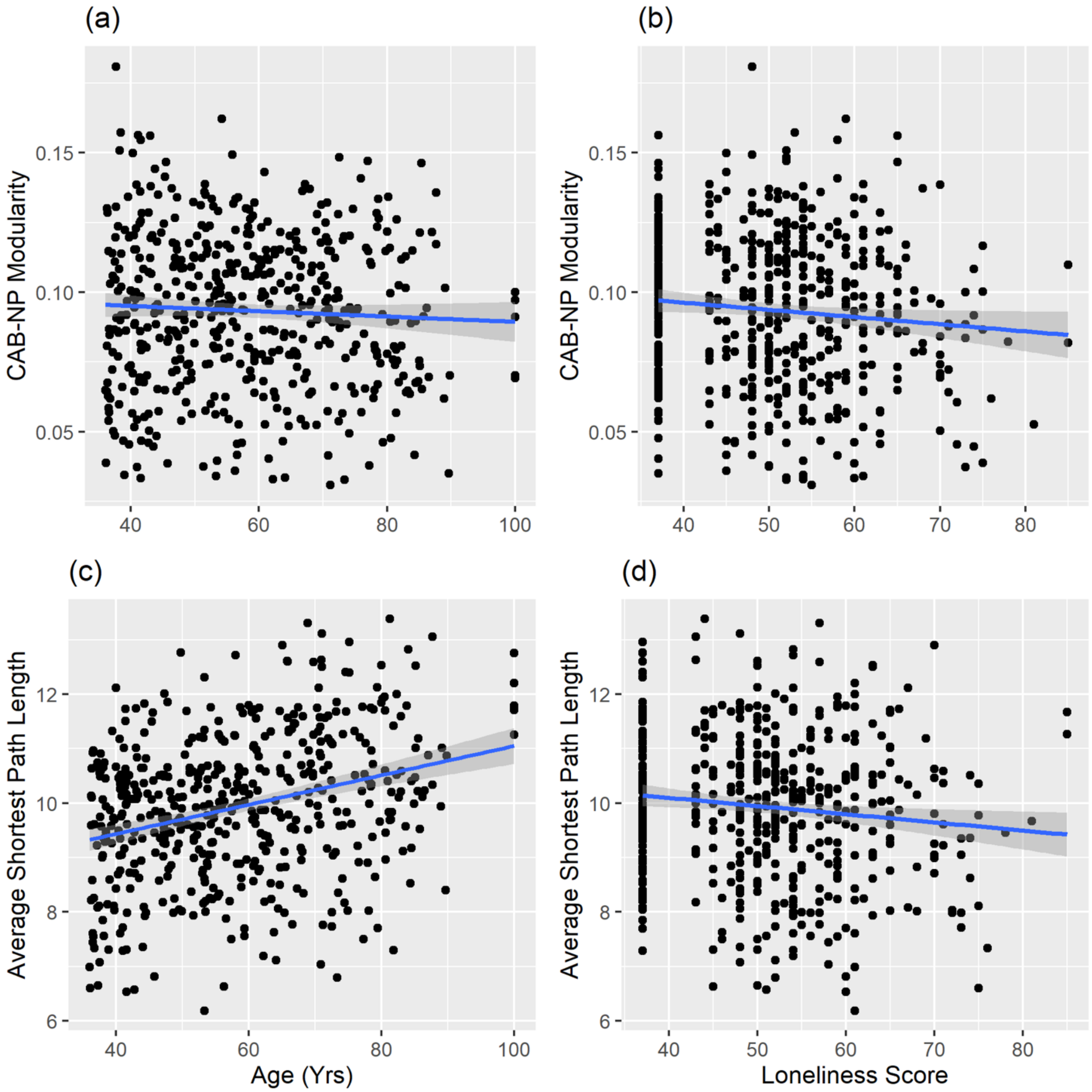

**Supplementary Figure S3**

*Age and Graph-Based Brain Functional Connectivity Association Maps: Negative-Only Connectivity Weights. Color bar shows the z-score from Aspin-Welch's test for using exchangeability blocks to adjusted for four scanning sites and variance was estimated for each block, implemented in FSL PALM.*

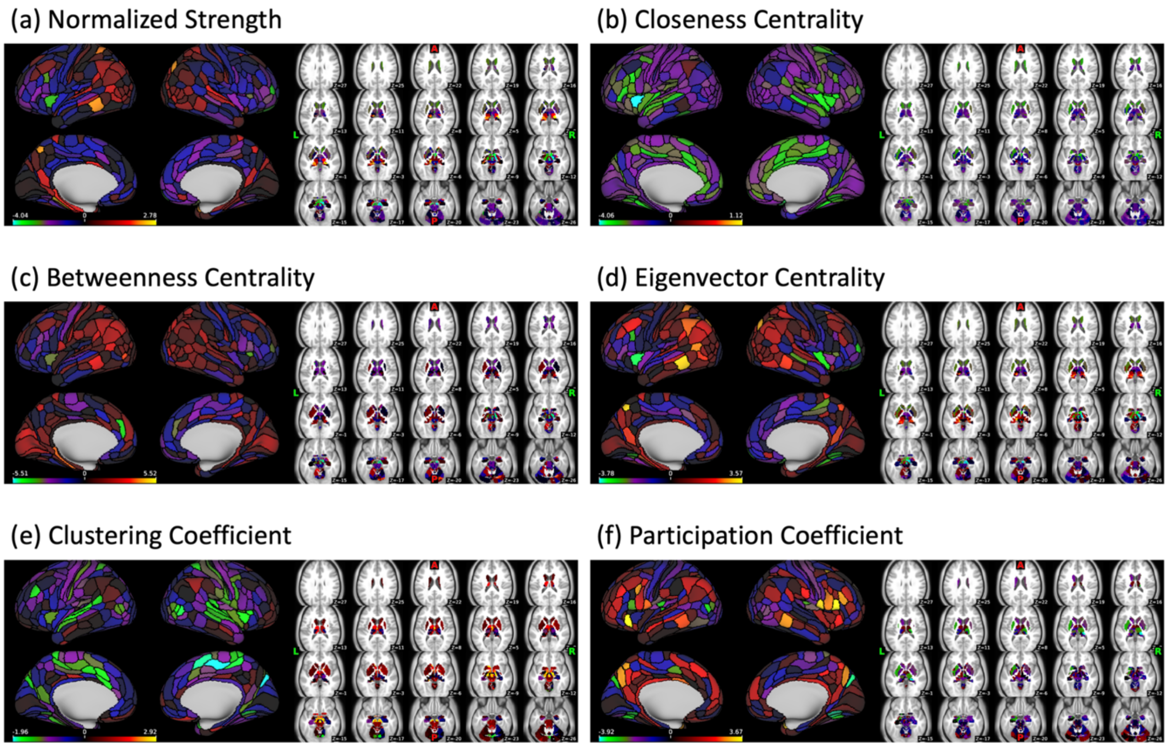

**Supplementary Figure S4**

*Association of Age and Global Graph-Based Brain Functional Connectivity: Negative-Only Connectivity Weights. Modularity association with (a) age and (b) loneliness. Average shortest path length with (c) age and (d) loneliness.*

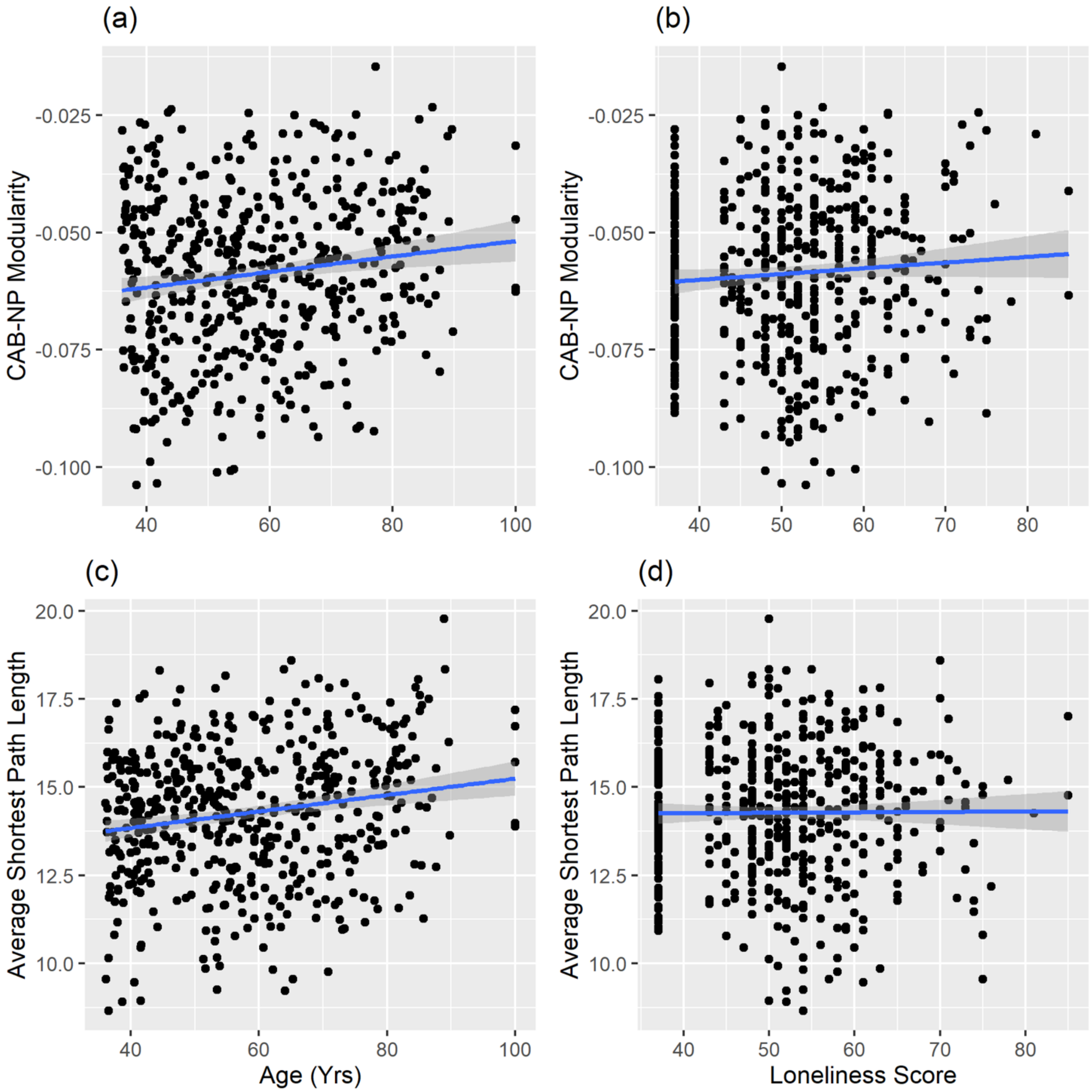

**Supplementary Figure S5**

*Figure is shown for illustrative purposes, depicting age-related trajectories in the loneliest (top tertile) and most socially connected (bottom tertile) individuals. The formal test of the loneliness × age interaction was conducted in PALM across the full sample (N = 512) using loneliness as a continuous variable.*

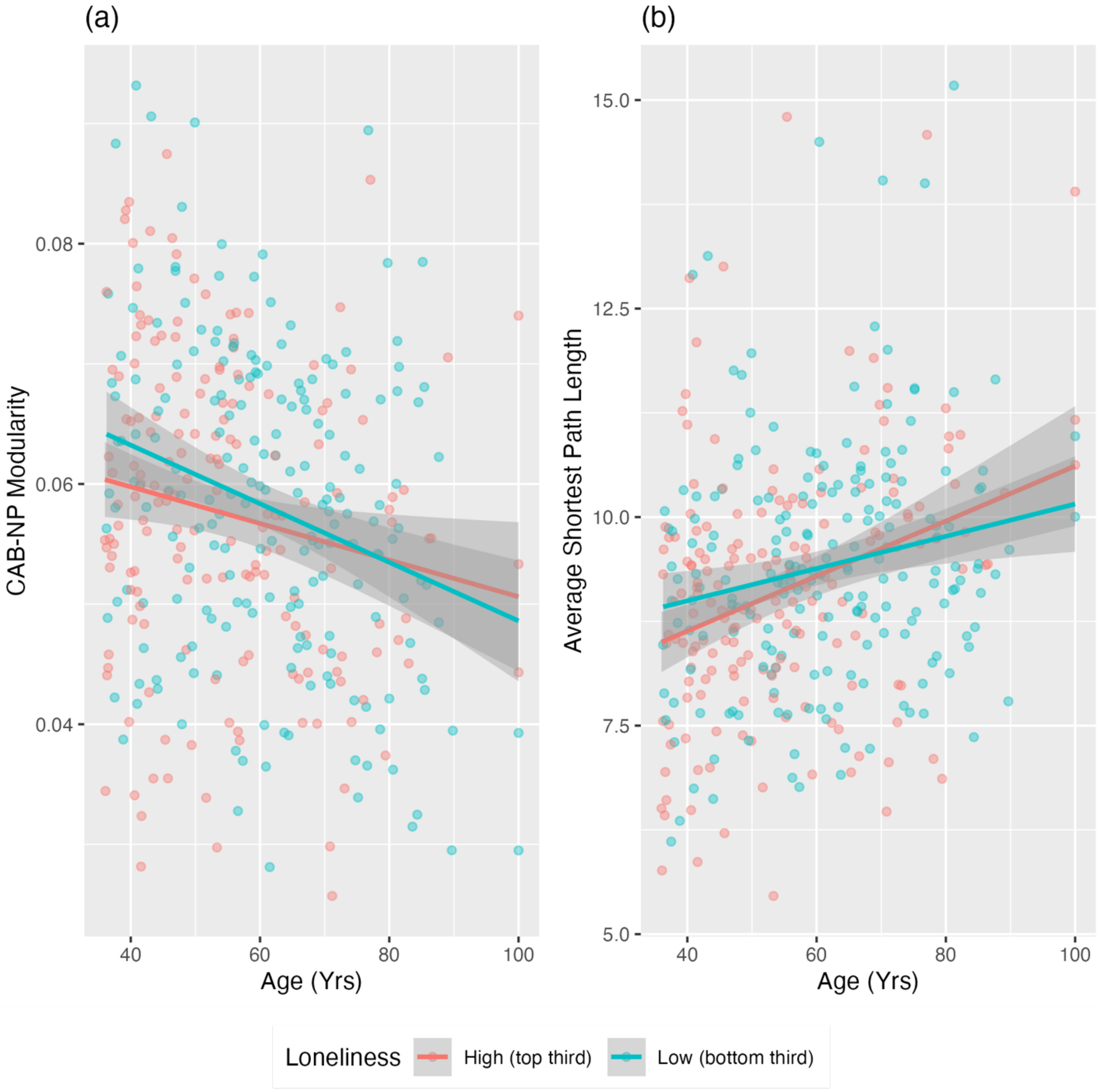
